# CTCF-mediated 3D chromatin predetermines the gene expression program in the male germline

**DOI:** 10.1101/2023.11.30.569508

**Authors:** Yuka Kitamura, Kazuki Takahashi, So Maezawa, Yasuhisa Munakata, Akihiko Sakashita, Noam Kaplan, Satoshi H. Namekawa

**Affiliations:** Department of Microbiology and Molecular Genetics, University of California, Davis, Davis, CA, 95616, USA; Division of Reproductive Sciences, Division of Developmental Biology, Perinatal Institute, Cincinnati Children’s Hospital Medical Center, Cincinnati, OH, 45229, USA; Faculty of Science and Technology, Department of Applied Biological Science, Tokyo University of Science, Noda, Chiba, 281-8510, Japan; Department of Molecular Biology, Keio University School of Medicine, Tokyo, 160-8582 Japan; Department of Physiology, Biophysics & Systems Biology, Rappaport Faculty of Medicine, Technion - Israel Institute of Technology, Haifa, Israel

## Abstract

Spermatogenesis is a unidirectional differentiation process that generates haploid sperm, but how the gene expression program that directs this process is established is largely unknown. Here we determine the high-resolution 3D chromatin architecture of male germ cells during spermatogenesis and show that CTCF-mediated 3D chromatin predetermines the gene expression program required for spermatogenesis. In undifferentiated spermatogonia, CTCF-mediated chromatin contacts on autosomes pre-establish meiosis-specific super-enhancers (SE). These meiotic SE recruit the master transcription factor A-MYB in meiotic spermatocytes, which strengthens their 3D contacts and instructs a burst of meiotic gene expression. We also find that at the mitosis-to-meiosis transition, the germline-specific Polycomb protein SCML2 resolves chromatin loops that are specific to mitotic spermatogonia. Moreover, SCML2 and A-MYB establish the unique 3D chromatin organization of sex chromosomes during meiotic sex chromosome inactivation. We propose that CTCF-mediated 3D chromatin organization enforces epigenetic priming that directs unidirectional differentiation, thereby determining the cellular identity of the male germline.

## Introduction

Eukaryotic genomes are folded into a dynamic three-dimensional (3D) architecture within the nucleus that influences gene expression^1–4^. The development of genome-wide chromosome conformation capture methods, especially Hi-C^5^, has accelerated our understanding of the 3D genome and the interplay between 3D genome organization and cell-fate decisions^6^. For example, disruption of 3D chromatin architecture has been shown to lead to disturbed gene expression, incomplete cell differentiation, and conversion to other cell types, at least in cell culture systems^7, 8^. Still, there is a major gap in our understanding of how the 3D genome defines gene expression programs during development and differentiation in vivo.

The mammalian male germline provides an ideal model to decipher the relationship between the 3D genome and gene expression programs. In spermatogenesis, after sex determination, male germ cells undergo a unidirectional differentiation process that comprises the maintenance of spermatogonia stem cells, commitment to meiosis, and production of haploid sperm^9^. Male germ cell differentiation is defined by chromatin-based mechanisms that instruct stage-specific gene expression both on autosomes and on sex chromosomes^10–12^. In spermatogonia, histone modifications are preset to regulate later gene expression programs^13–16^. Specifically, on autosomes, dimethylation of histone H3 at lysine 4 (H3K4me2) is pre-established on meiotic super-enhancer (SE) loci that drive a genome-wide burst of transcription after the mitosis-to-meiosis transition^17^. The sex chromosomes, on the other hand, undergo meiotic sex chromosome inactivation (MSCI), an essential event in the male germline^11, 18^. They form a distinct nuclear compartment called the XY body (sex body) that is physically segregated from the autosomes^19^.

In this study, we elucidated how the 3D genome architecture of male germ cells is regulated to define the gene expression programs that drive spermatogenesis. Drawing on the recent analysis of basic 3D chromatin features in spermatogenesis^20–24^, we performed high-resolution Hi-C analysis using cell types representative of major stages of spermatogenesis to decipher the detailed pictures of the 3D genome in the male germline at unprecedented resolution. To determine how the 3D genome regulates the dynamic transcriptional transition from the mitotic to meiotic stages, we also performed Hi-C analyses using mouse mutants lacking key transcriptional regulators of spermatogenesis. One of these factors is SCML2, a germ cell-specific component of Polycomb Repressive Complex 1 (PRC1) that is critical for the suppression of the mitotic program in late spermatogenesis^2^. The other is A-MYB (MYBL1), a master transcription factor that regulates the burst of meiotic gene expression at the pachytene stage^25^. These functional analyses reveal that SCML2 resolves the mitotic 3D chromatin organizatiton, whereas A-MYB drives the establishment of meiotic 3D chromatin. Importantly, we show that the unidirectional differentiation program during spermatogenesis is predetermined by CTCF-mediated 3D chromatin contacts. These results provide a molecular basis for the cellular identity of male germ cells defined by the 3D genome.

## Results

### High-resolution Hi-C data sets reveal 3D chromatin reprograming during spermatogenesis

To determine high-resolution 3D chromatin structures of germ cells during the course of spermatogenesis, we performed in-depth Hi-C analysis of spermatogenic cells isolated at four representative developmental stages (Fig. 1a). Specifically, we isolated THY1^+^ undifferentiated spermatogonia and KIT^+^ differentiating spermatogonia from the testes of 7-day-old male C57BL/6 mice using magnetic-activated cell sorting (MACS)^26^. In addition, we isolated pachytene spermatocytes (PS), which are in the meiotic prophase, and round spermatids (RS), which are in the postmeiotic stage, from the testes of adult male C57BL/6 mice using BSA gradient sedimentation^20^. We confirmed high purity for all cell types (Extended Data Fig. 1a, b) and performed Hi-C experiments on two biological replicates for each cell type (Supplementary Table 1). These replicates showed a high correlation (Extended Data Fig. 1c) and we merged them for downstream analysis, yielding ∼430-670 million Hi-C contact reads for each stage and exceeding the total read depths of previous Hi-C analyses in spermatogenesis^20–24^. Comparing the four developmental stages, we detected a relatively high correlation between mitotic spermatogonia (THY1^+^ and KIT^+^) compared to PS and RS (Extended Data Fig. 1d). This reflects the biological similarity between THY1^+^ and KIT^+^ spermatogonia and the presence of a dynamic transition from the mitotic stages to the meiotic and postmeiotic stages.

**Figure 1.**
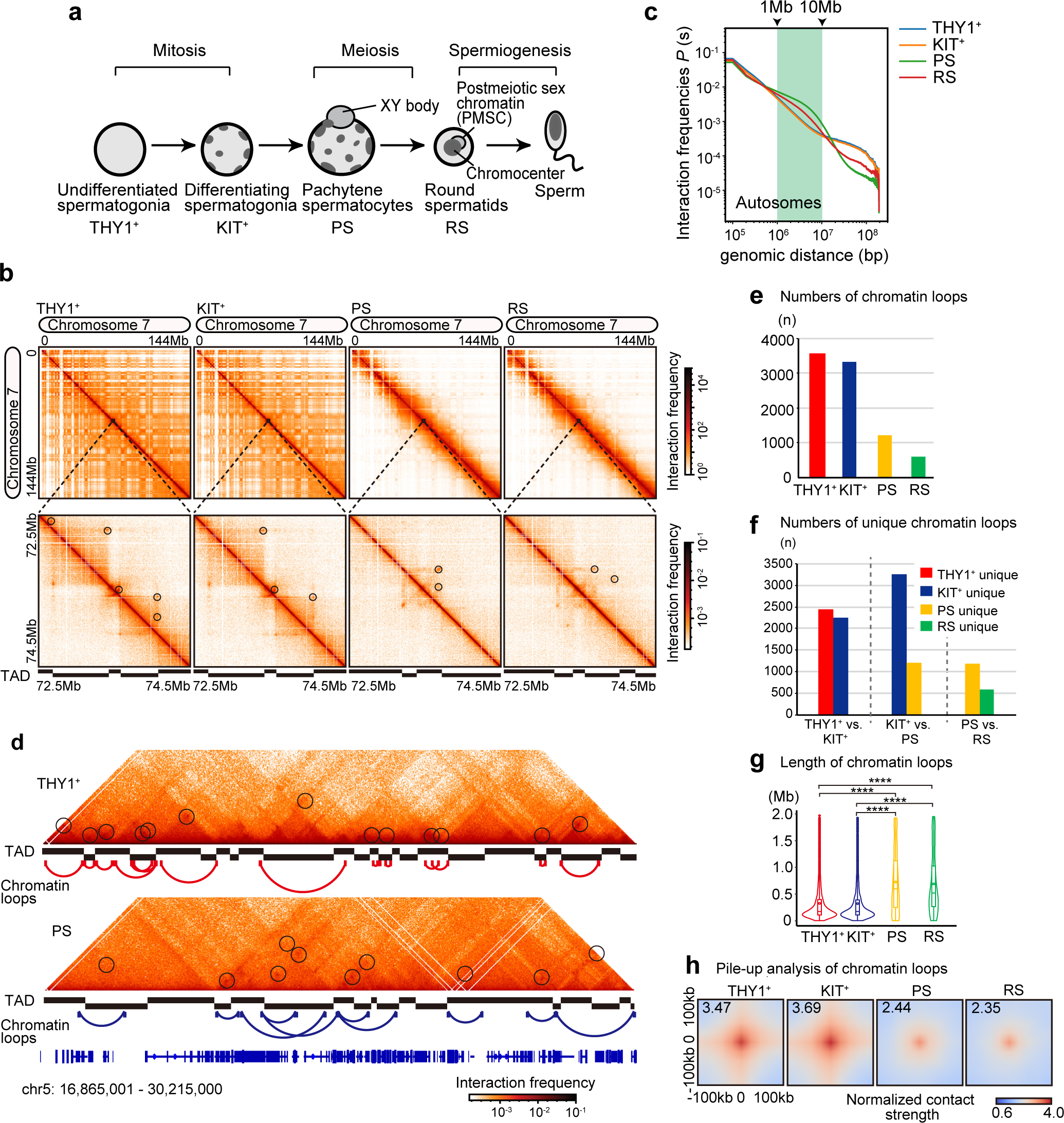
3D chromatin reprogramming and inter-TAD chromatin loop formation in meiosis. **a**, Schematic of the stages of mouse spermatogenesis analyzed in this study. THY1^+^: undifferentiated spermatogonia; KIT^+^: differentiating spermatogonia; PS: pachytene spermatocytes; RS: round spermatids. **b**, Hi-C maps showing normalized Hi-C interaction frequencies (100kb bins, chromosome 7) in THY1^+^, KIT^+^, PS, and RS. 10kb bins normalized Hi-C matrices were used for the zoom-in. Black circles in the Hi-C map indicate chromatin loops. **c**, Hi-C interaction frequency probabilities *P* stratified by genomic distance *s* for each cell type shown (100kb bins). All autosomes were analyzed. **d**, Hi-C interaction heat maps (25kb bins, chromosome 5, 16, 865, 001-30, 215, 000bp) in THY1^+^ and PS. Chromatin loops are indicated by black circles, red lines in THY1^+^, and blue lines in PS. **e**, Numbers of chromatin loops (n) detected from each Hi-C data set (merged results for each using 5kb, 10kb, and 25kb bin data). **f**, Numbers of unique chromatin loops comparing each pairwise developmental stage. **g**, Chromatin loop length (Mb) from each Hi-C data set (merged results for each using 5kb, 10kb, and 25kb bin data). The number of loops used in the analysis was equal to the number shown in e (THY1^+^: n=3,562, KIT^+^: n=3,336, PS: n=1,223, RS: n=609). The box indicates the 25th, median and 75th percentiles, and the dot in the box indicates mean. Statistical analysis is based on Bonferroni correction. **** indicates *p* < 2e^-16^. **h**, Chromatin loop pile-up in each cell type with 100kb padding. Color represents normalized contact strength in the log scale. The normalized contact strength values in the central pixel are shown on the top left.

Next, we examined the Hi-C maps of an entire representative chromosome (chromosome 7) at each stage, including a zoom-in to a specific chromosomal region (Fig. 1b). Previous high-resolution Hi-C studies in other cellular systems revealed the presence of point interactions that represent stable chromatin-loops (dots) in addition to topologically associating domains (TADs), which manifest in triangular patterns^27–29^. We also detected these stable chromatin loops (“chromatin loops” hereafter) (Fig. 1b, bottom, marked by circles), confirming the high resolution of our new data sets. The Hi-C interaction contact matrices show that the genomes of THY1^+^ and KIT^+^ cells are enriched in distal interactions (>10 Mb), a typical feature of interphase nuclei. These interactions are abolished in PS, where proximal interactions (a range between 1-10 Mb) dominate (Fig. 1b). This tendency was confirmed by a contact probability *P(s)* analysis, which is indicative of the general polymer state of chromatin^5, 30^ (Fig. 1c). These results corroborate previous Hi-C studies^21, 22^ and confirm that the typical interphase pattern of high-order chromatin present in spermatogonia is reprogrammed when cells enter meiosis.

### Formation of inter-TAD chromatin loops for meiotic gene regulation

Chromatin is spatially organized into TADs, which restrict interactions of cis-regulatory sequences and thereby contribute to gene regulation^31–33^. Accordingly, chromatin loops, which are critical for gene expression regulation^1, 34, 35^, are typically observed within TADs. Indeed, in THY1^+^ spermatogonia, chromatin loops were largely detected within TADs (intra-TADs: Fig. 1d). By contrast, in PS, many chromatin loops were detected beyond TAD borders (inter-TADs, Fig. 1d). From mitotic spermatogonia to meiotic PS, the total number of chromatin loops decreased (Fig. 1e). Of note, the chromatin loops present in THY1^+^ spermatogonia and PS are mostly unique, and the same holds true for the subsequent PS to RS transition (Fig. 1f). This feature presumably reflects formation and resolution of meiotic chromosome structure, organized into chromatin loop arrays along chromosome axes. Moreover, the length of chromatin loops increased from mitotic spermatogonia to meiotic PS (Fig. 1g), while average contact strengths decreased based on a pile-up analysis^36^ (Fig. 1h). These results demonstrate that the structural changes that occur at the mitosis to meiosis transition are based on the resolution of intra-TAD chromatin loops and the de novo establishment of inter-TAD chromatin loops.

The weakening of TADs in PS^20–24^ indicates that this feature might drive meiotic gene regulation mediated by inter-TAD chromatin loops. To test this possibility, we investigated the relationship between TAD strength and chromatin loop formation during spermatogenesis. First, we detected TADs in each stage of spermatogenesis. The number of TADs decreased during the transition from spermatogonia to meiotic PS and recovered in postmeiotic RS (Fig. 2a). This is consistent with previous reports showing the attenuation of TADs in meiotic prophase^21, 22^. TAD boundaries were largely shared between THY1^+^ and KIT^+^ spermatogonia, with 3,550 common TAD boundaries (85% of 4,192 THY1^+^ TAD boundaries), and they progressively changed from KIT^+^ to PS and from PS to RS (Fig. 2b). Among 4,039 TAD boundaries in KIT^+^ spermatogonia, only 1,447 (36%) were maintained in PSs. In contrast, from PSs to RSs, 1,935 (80%) out of 2,420 PS TAD boundaries were maintained, and 1,659 TAD boundaries were newly generated in RS (Fig. 2b). To define stage-specific features of TADs, we next examined the average contact strength of TADs using a pile-up analysis of the Hi-C matrices, which visualizes average insulation strengths of the regions around TADs and their boundaries^36^. We found that while insulation at TAD boundaries was weakened during the transition from KIT^+^ spermatogonia to PS (Fig. 2c), the interaction strengths between adjacent TADs increased (Fig. 2d). Thus, during the transition from mitotic spermatogonia to meiotic PS, TADs and TAD borders are reprogrammed. Interactions beyond TAD boundaries increase in meiosis, not only through the de novo formation of inter-TAD chromatin loops but also through TAD-TAD interactions.

**Figure 2.**
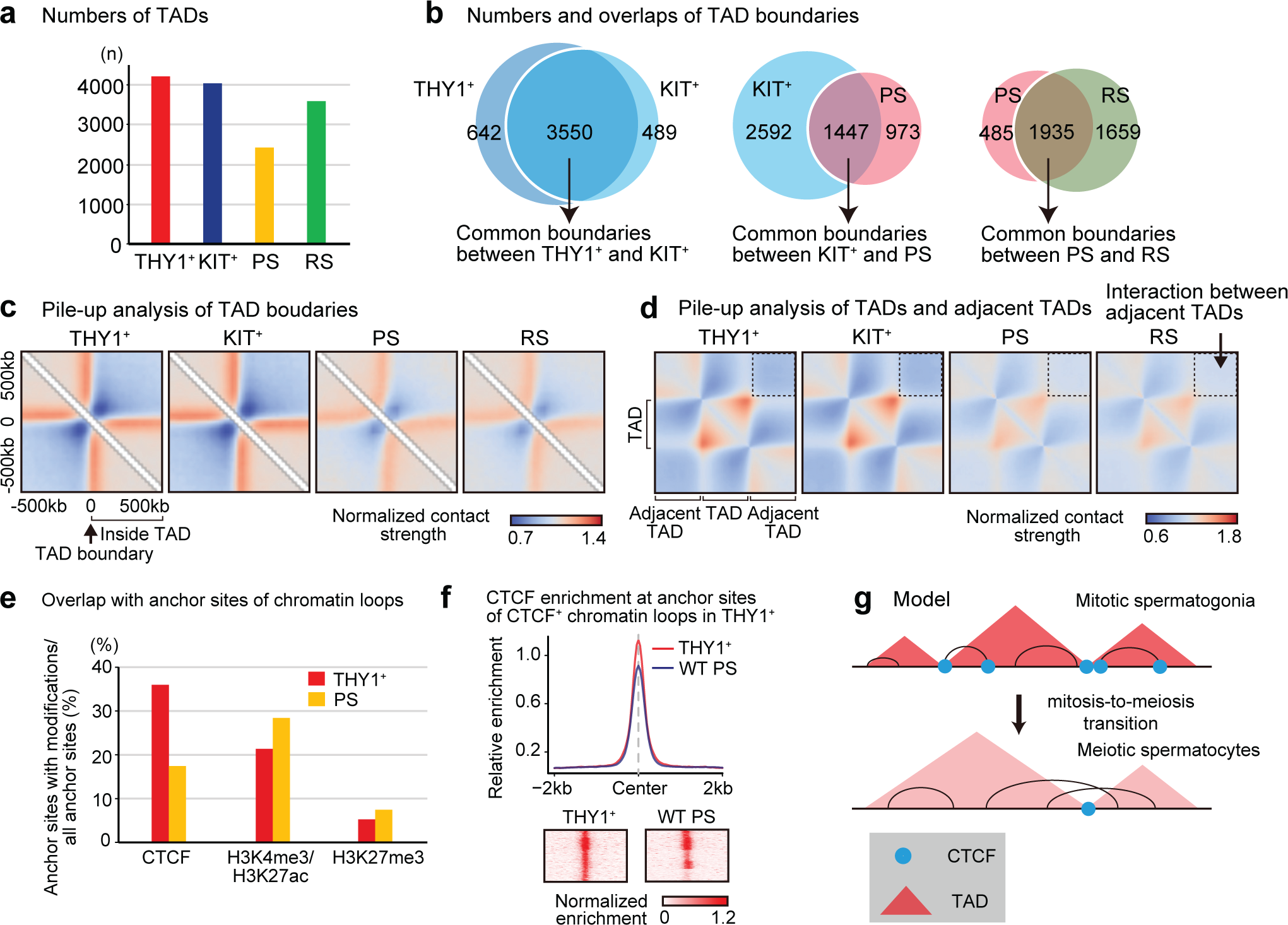
TAD and chromatin loop reorganization during spermatogenesis. **a**, Numbers of TADs (n) detected from each Hi-C data set (25kb bins). **b**, Venn diagram showing numbers and overlaps of TAD boundaries in each developmental stage. **c**, Local pile-up analysis of TAD boundaries in each cell type. 10kb bins Hi-C data with 500 kb padding around the central pixel. Color represents normalized contact strength in the log scale. **d**, Local rescaled pile-ups of TADs from 10kb bin Hi-C data in each cell type. The dotted regions represent interactions between adjacent TADs. **e**, Ratio of accumulation of CTCF, H3K4me3/H3K27ac, or H3K27me3 at the anchor sites of chromatin loops in THY1^+^ and PS. **f**, CTCF enrichment at anchor sites of CTCF-dependent chromatin loops in THY1^+^ (detected in panel **e**, 2,532 sites). Heat maps for each locus are shown at the bottom. **g**, Model showing TAD and chromatin loop reorganization at the mitosis-to-meiosis transition.

There are various types of chromatin loops, including CTCF-CTCF loops, enhancer-promoter loops, and Polycomb-dependent loops. Since these different types contribute to gene activation or silencing^28, 37^, it is possible to infer their functions. Therefore, we examined the modifications on anchor sites of chromatin loops and distinguished the three loop classes based on the presence of CTCF, H3K4me3 (a promoter mark)/H3K27ac (an active enhancer mark), or H3K27me3 (the Polycomb repressive complex2 (PRC2)-mediated mark). First, we performed CTCF ChIP-seq in THY1^+^ spermatogonia and PS (Extended Data Fig. 2a). CTCF binding sites overlapped with 35.6% of anchor sites of THY1^+^ chromatin loops, while this overlap decreased to 17.3 % in PS, suggesting the resolution of CTCF-pair loops during meiosis (Fig. 2e). Indeed, at some of the sites of CTCF-pair loops in THY1^+^ spermatogonia, CTCF enrichment was reduced or lost in PS (Fig. 2f). This suggests that the loss of CTCF might underly the resolution of some CTCF-pair loops in meiosis. Second, we used our previous H3K4me3, H3K27ac, and H3K27me3 ChIP-seq datasets in THY1^+^ and PSs^17, 38, 39^ and defined enhancer/promoter-pair loops and Polycomb-dependent loops. The proportion of enhancer/promoter-pair loops was in the range of ∼20-30 %, and the proportion of Polycomb-dependent loops was less than 10 % of total chromatin loops in THY1^+^ and PS (Fig. 2e). Since chromatin loops in PS are mostly PS-specific (Fig. 1f), all three types of chromatin loops are largely de novo generated during meiosis. Contact strengths were comparable within each class of loops in THY1^+^ and PS (Extended Data Fig. 2b), indicating that contact strength is not class-but stage-dependent.

We further examined how the resolution of TADs at the mitosis-to-meiosis transition is regulated. As opposed to the resolution of CTCF-pair loops (Fig. 2e, f), there was no change in the proportion of CTCF-associated TAD boundaries among all TAD boundaries (Extended Data Fig. 2c). This suggests that distinct mechanisms operate between the resolution of TAD boundaries and chromatin loops in meiosis. Taken together, these results demonstrate that attenuation of TADs and resolution of CTCF-pair loops take place at the mitosis-to-meiosis transition to establish long inter-TAD loops during meiosis (Fig. 2g).

### SCML2 is required for the resolution of spermatogonia-type 3D chromatin and gene repression

To further understand the mechanisms that underlie the resolution of TAD boundaries and chromatin loops at the transition from mitotic spermatogonia to meiotic spermatocytes, we focused on the germline-specific Polycomb protein SCML2. SCML2 is responsible for the suppression of genes that are highly expressed in mitotic spermatogonia after the mitosis-to-meiosis transition^14^. It is expressed in undifferentiated spermatogonia and forms part of PRC1, which deposits H2AK119ub^14^ and facilitates PRC2-mediated H3K27me3 during meiosis^39^. H3K27me3 counteracts the active enhancer mark H3K27ac, thereby resolving spermatogonia-type enhancers^17^. We therefore hypothesized that SCML2 is involved in the resolution of spermatogonia-type 3D chromatin. To test this hypothesis, we performed Hi-C analysis using *Scml2* knockout (*Scml2*-KO) PSs and RSs (Extended Data Fig. 1c). *Scml2*-KO PSs and RSs showed increased distal interactions compared to wild-type cells, and this pattern resembles Hi-C maps in THY1^+^ and KIT^+^ cells (Fig. 3a). Pearson correlation analysis also showed that *Scml2*-KO PSs and RSs are more similar to wild-type THY1^+^ and KIT^+^ than wild-type PSs and RSs (Extended Data Fig. 1d). This suggests that the mitotic 3D chromatin organization of spermatogonia is retained in *Scml2*-KO cells during meiosis.

**Figure 3.**
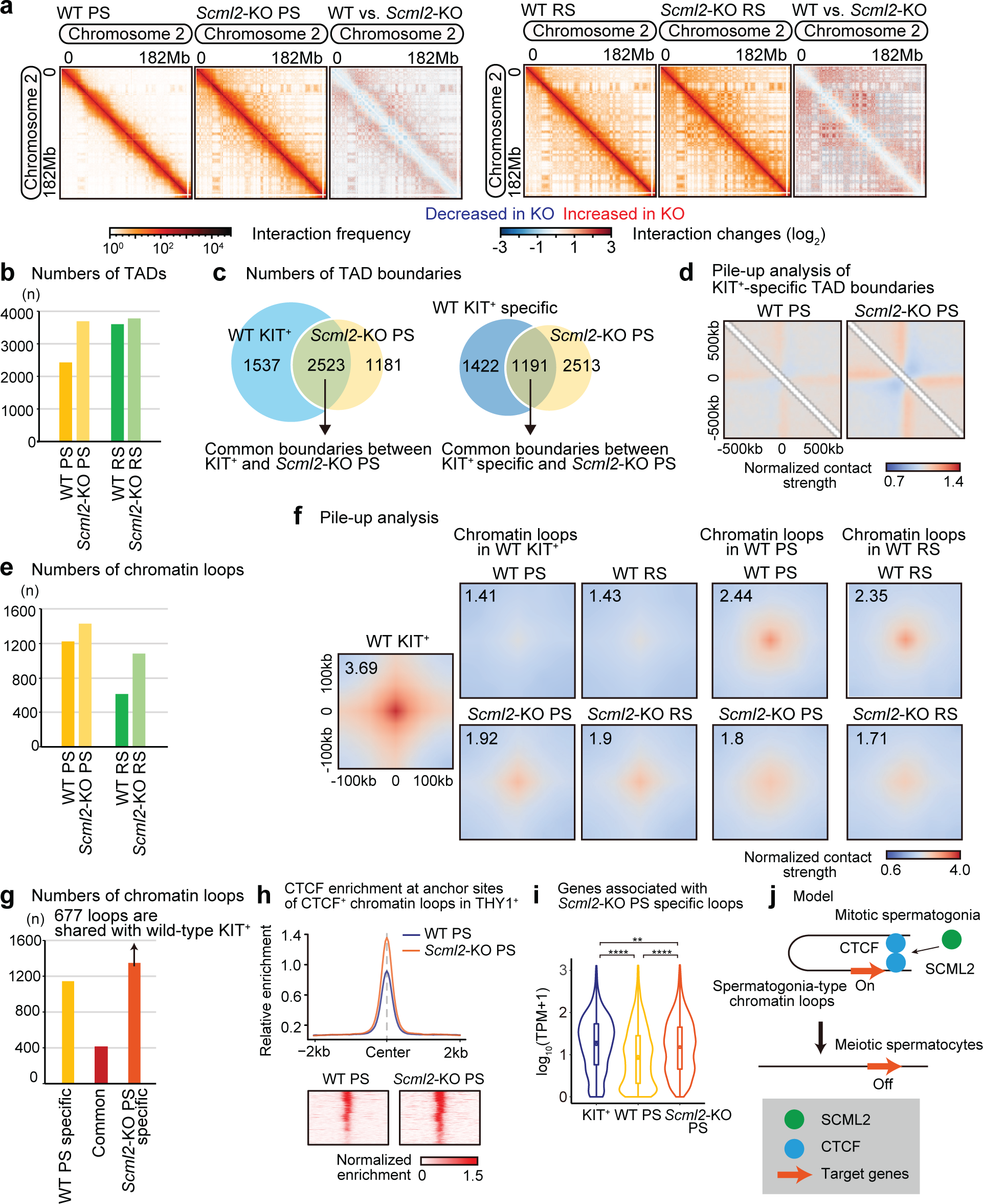
SCML2 is required for the resolution of spermatogonia-type 3D chromatin. **a**, Heat maps showing normalized Hi-C interaction frequencies (100kb bins, chromosome 2) in wildtype (WT) PS, *Scml2*-KO PS (left), and WT RS and *Scml2*-KO RS (right). Red and blue Hi-C maps represent a log2 ratio comparison of Hi-C interaction frequencies between WT and *Scml2*-KO. **b**, Numbers of TADs (n) detected from each Hi-C data set (25kb bins) in WT PS, *Scml2*-KO PS, WT RS, and *Scml2*-KO RS. **c**, Venn diagram showing the overlap between all KIT^+^ TAD boundaries and *Scml2*-KO PS TAD boundaries (left), and the overlap between KIT^+^-specific TAD boundaries and *Scml2*-KO PS TAD boundaries (right). KIT^+^-specific boundaries are defined by excluding TAD boundaries detected in WT PS. **d**, Local pile-up analysis of KIT^+^ specific TAD boundaries in WT PS and *Scml2*-KO PS. **e**, Numbers of chromatin loops (n) detected in each Hi-C data set (merged results for each using 5kb, 10kb, and 25kb bin data) in WT PS, *Scml2*-KO PS, WT RS, and *Scml2*-KO RS. **f**, Chromatin loop pile-up analysis in each cell type with 100kb padding. The normalized contact strength in the central pixel is displayed on the top left. **g**, Numbers of specific and common chromatin loops between WT PS and *Scml2*-KO PS. 677 *Scml2*-KO PS-specific loops overlapped with loops detected in KIT^+^. Overlapping loops were detected by Juicer. **h**, CTCF enrichment in WT PS and *Scml2*-KO PS at the anchor site of CTCF-chromatin loops in THY1^+^ spermatogonia. **i**, Violin plots of RNA-seq reads converted to log10 (TPM+1) value for genes associated with *Scml2*-KO PS specific loops in KIT^+^, WT PS and *Scml2*-KO PS. 1,243 genes were identified by extracting genes present in the anchor site of *Scml2*-KO PS-specific loops. The box indicates the 25th, median and 75th percentiles, and the dot in the box indicates mean. Statistical analysis is based on Bonferroni correction. ****: *p* < 2e^-16^, **: *p* < 0.005. **j**, Model of resolution of spermatogonia-type 3D chromatin by SCML2.

To determine whether SCML2 mediates the resolution of spermatogonia-type TAD boundaries, we detected TADs in the Hi-C dataset of *Scml2*-KO PS and RS. The number of TADs increased in *Scml2*-KO PS compared to wild-type PS (Fig. 3b). Of note, TAD boundaries in the *Scml2*-KO PSs overlapped with the TAD boundaries in wild-type KIT^+^ spermatogonia at 2,523 loci (Fig. 3c). This overlap is more abundant than that between KIT^+^ and wild-type PS TAD boundaries (1,447 loci; Fig. 3c), suggesting that spermatogonia-type TAD boundaries are retained in *Scml2*-KO PS. Indeed, among 2,613 KIT^+^-specific TAD boundaries (which do not overlap with wild-type PS TAD boundaries), 1,191 loci remain in *Scml2*-KO PS TAD boundaries. Pile-up analysis further confirmed that KIT^+^-specific TAD boundaries remain in *Scml2*-KO PS (Fig. 3d). These results indicate that SCML2 is required for the resolution of KIT^+^-specific TADs in PS.

Next, we examined the role of SCML2 in the resolution of chromatin loops. The number of chromatin loops increased in *Scml2*-KO PSs and RSs compared to wild-type PS and RS (Fig. 3e). While wild-type KIT^+^-specific chromatin loops did not show high contact strengths in wild-type PS and RS, they remained in *Scml2*-KO PS and RS (Fig. 3f). On the other hand, chromatin loops detected in wild-type PS and RS did not show high contact strengths in *Scml2* KO PS and RS (Fig. 3f). Comparison of chromatin loops between wild-type PSs and *Scml2*-KO PSs revealed that there were 1,358 *Scml2*-KO-specific chromatin loops, 677 of which are shared with chromatin loops present in wild-type KIT^+^ (Fig. 3g). The persistence of KIT^+^ chromatin loops in *Scml2* KO PS and RS was confirmed with Hi-C maps (Extended Data Fig. 3a). Therefore, SCML2 is also involved in the resolution of spermatogonia-type chromatin loops.

To determine how SCML2 resolves spermatogonia-type 3D chromatin, we next examined whether SCML2 is required for the resolution of CTCF sites by performing CTCF ChIP-seq in *Scml2*-KO PS. The Pearson correlation between *Scml2*-KO PS and wild-type THY1^+^ spermatogonia (0.79) is higher than the Pearson correlation between wild-type PS and wild-type THY1^+^ (0.73; Extended Data Fig. 3c), suggesting that CTCF distribution in *Scml2*-KO PS is more similar to wild-type THY1^+^ than that in wild-type PS. In wild-type PS, CTCF enrichment at the anchor sites of the CTCF pair loops detected in THY1^+^ was reduced (Fig. 2f), but CTCF enrichment at these loci remained high in *Scml2*-KO PS (Fig. 3h). This suggests that SCML2 is involved in the resolution of at least a fraction of CTCF sites.

Since chromatin conformation is implicated in the regulation of gene expression^31, 33^, we examined the effect of the KIT^+^-specific loops that persist in *Scml2*-KO PS on gene expression. Therefore, we examined the expression profile of 1,243 genes present in the anchor sites of the *Scml2*-KO PS-specific loops defined in Fig. 3g. The overall expression level of these genes decreased from wild-type KIT^+^ to wild-type PSs, but remained high in *Scml2*-KO PS compared to wild-type PSs (Fig. 3i). We thus conclude that SCML2 is required for the resolution of spermatogonia-type 3D chromatin, thereby suppressing spermatogonia-type gene expression in meiosis (Fig. 3j). Importantly, we did not observe a significant change in contact strengths of chromatin loops at SCML2-dependent bivalent promoters marked by both active (H3K4me2/3) and repressive (H3K27me3) histone modifications (Extended Data Fig. 3d). Therefore, the function of SCML2 in resolving spermatogonia-type 3D chromatin is independent of its regulation of bivalent promoters^39^.

### A-MYB is required for the formation of meiotic-type 3D chromatin and gene activation

Because meiosis-specific chromatin loops are *de novo* generated after the resolution of spermatogonia-type chromatin loops (Fig. 2g), we next sought to determine the mechanism driving meiosis-specific chromatin loops. To this end, we focused on A-MYB, a transcription factor responsible for the activation of pachytene-specific genes^25^. A-MYB is required to establish H3K27ac on pachytene-specific enhancers, thereby activating these enhancers^17^. We suspected a role of A-MYB in the formation of meiosis-specific chromatin loops because of the establishment of specific enhancer/promoter-pair loops in PS. We therefore isolated PS from *A-myb* mutant (*Mybl1^repro9^*) mice and performed Hi-C analysis. We found that distal interactions were increased in the *A-myb* mutant PS as shown in a Hi-C heat map (Fig. 4a) and in a contact probability analysis (Extended Data Fig. 4a), suggesting that spermatogonia-type 3D chromatin is retained in the *A-myb* mutant PS. In accordance with this notion, the number of TADs also increased (Fig. 4b); in fact, more than 70% of the TAD boundaries in the *A-Myb* mutant PS were common to those detected in KIT^+^ (Fig. 4c: left), and KIT^+^-specific loops largely remained in the *A-myb* mutant PS (Fig. 4c: right). Further, KIT-specific TAD boundaries retained high contact strength in the *A-myb* mutant PS (Fig. 4d). These results demonstrate that A-MYB is required for the establishment of meiosis-type 3D chromatin, and that its loss leads to the retention of spermatogonia-type 3D chromatin.

**Figure 4.**
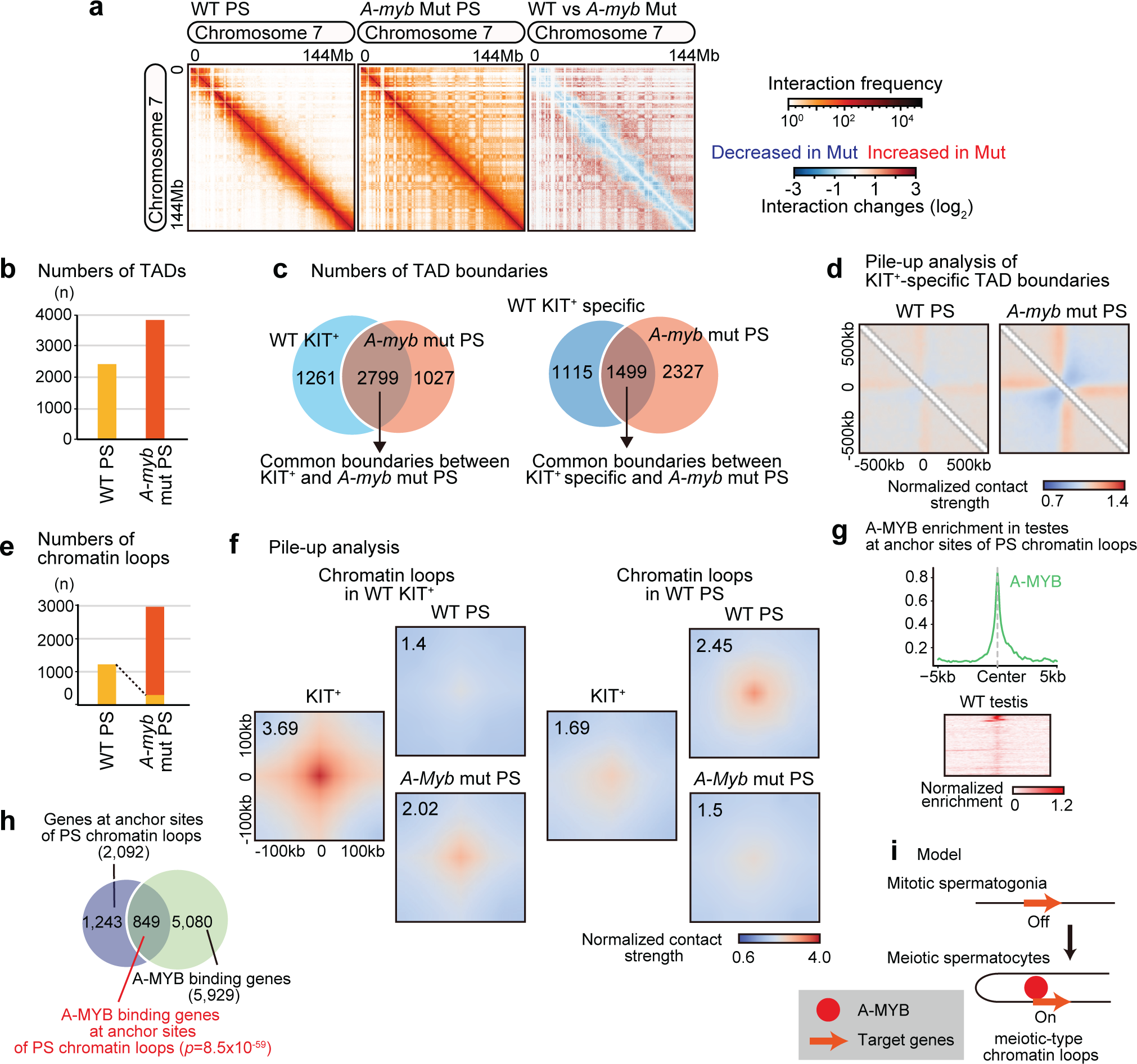
A-MYB is required for the formation of meiotic-type 3D chromatin. **a**, Heat maps showing normalized Hi-C interaction frequencies (100kb bins, chromosome 7) in WT PS, *A-myb* mutant PS (left). Red and blue Hi-C maps represent a log2 ratio comparison of Hi-C interaction frequencies between wild-type and *A-myb* mutant PS. **b**, Number of TADs (n) detected from each Hi-C data set (25kb bins) in WT PS and *A-myb* mutant PS. **c**, Venn diagram showing the overlap between all KIT^+^ TAD boundaries and *A-myb* mutant PS TAD boundaries (left), and the overlap between KIT^+^-specific TAD boundaries and *A-myb* mutant PS TAD boundaries (right). KIT^+^-specific boundaries are defined by excluding boundaries detected in WT PS. **d**, Local pile-up analysis of KIT^+^-specific TAD boundaries in WT PS and *A-myb* mutant PS. **e**, Number of chromatin loops (n) detected in each Hi-C data set (merged results for each using 5kb, 10kb, and 25kb bin data) in WT PS and *A-myb* mutant PS. Yellow area in the graph of *A-myb* mutant PS indicate that the same loops are detected in WT PS (357 loops). **f**, Chromatin loop pile-up analysis in each cell type with 100kb padding. The normalized contact strength in the central pixel is displayed on the top left. **g**, ChIP-seq data for A-MYB using whole testis at the regions adjacent to TSS of 849 genes that overlap with anchor sites of chromatin loops in PS. **h**, Venn diagram showing the intersection of genes located at anchor sites of chromatin loops in PS (blue) and all A-MYB bound genes (green). The overlap is statistically significant (*p*=8.5x10^-59^) compared to the proportion of all A-MYB bound genes to all RefSeq genes based on the hypergeometric test. **i**, Model of the establishment of meiotic-type chromatin loops by A-MYB.

We next examined whether A-MYB is required to establish meiosis-specific chromatin loops. Although the total number of chromatin loops increased in the *A-myb* mutant PS compared to wild-type PS, only 29 % of wild-type PS-specific chromatin loops (357 out of 1,223) were detected in the *A-myb* mutant PS. Pile-up analyses show that *A-myb* mutant PS retained the contact strength of KIT^+^ chromatin loops (Fig. 4f, left), while *A-myb* mutant PS did not show high contact strength for PS chromatin loops (Fig. 4f). To test whether A-MYB directly mediates the formation of chromatin loops in PS, we reanalyzed previous ChIP-seq data of A-MYB using whole testis^25^. A-MYB is enriched at the anchor sites of PS chromatin loops (Fig. 4g), and A-MYB binds to 41% of genes at anchor sites of PS (849 out of 2,092: Fig. 4h). This association is statistically significant when compared to the ratio of all A-MYB binding genes to all RefSeq genes (5,929/22,661; P = 8.5×10^-59^, Hypergeometric test). These results indicate that A-MYB mediates the formation of a large part of chromatin loops in PS (Fig. 4i).

### A-MYB-dependent 3D chromatin is associated with the production of pachytene piRNAs

Another major function of A-MYB is the production of pachytene piRNAs^40^, which are involved in the maintenance of genome integrity and gene regulation in late spermatogenesis^41, 42^. A-MYB drives the production of pachytene piRNAs in a parallel mechanism with its regulation of enhancers through the induction of H3K27ac at pachytene piRNA clusters^17^. The loci of pachytene piRNA clusters switch from the B compartment to the A compartment during the mitosis-to-meiosis transition^22^. Our Hi-C data showed that 3D chromatin contacts were specifically detected at pachytene piRNA clusters in PS and were retained in RS, and that the formation of 3D chromatin is A-MYB dependent (Extended Data Fig. 4b). Thus, the A-MYB-mediated formation of 3D chromatin is associated with the production of pachytene piRNAs.

### Meiotic super-enhancers are poised within 3D chromatin

We next sought to determine how the global transcriptional changes that occur during spermatogenesis are regulated in the context of the 3D genome. To address this question, we focused on super-enhancers (SEs), which are long stretches of enhancers that play a central role in driving cell-type-specific gene expression and determining cellular identities^43–45^. In PS, A-MYB activates meiosis-specific SEs (meiotic SEs) through the establishment of H3K27ac to drive the expression of late spermatogenesis-specific genes^17^. Based on the specific enrichment of H3K27ac, we defined 399 meiotic SEs on autosomes, which are specific to PS. Among these, 270 (67.7%) are associated with PS chromatin loops (Extended Data Fig. 5a). A chromosome-wide track view confirms that chromatin loops (detected as stable chromatin loops in this study) are largely associated with SEs and the active genic loci that are in context with SEs (Fig. 5a). Therefore, these stable chromatin loops are associated with gene regulation and are distinct from meiotic chromatin loop arrays that are formed along the chromosome axes during meiotic prophase^46^.

**Figure 5.**
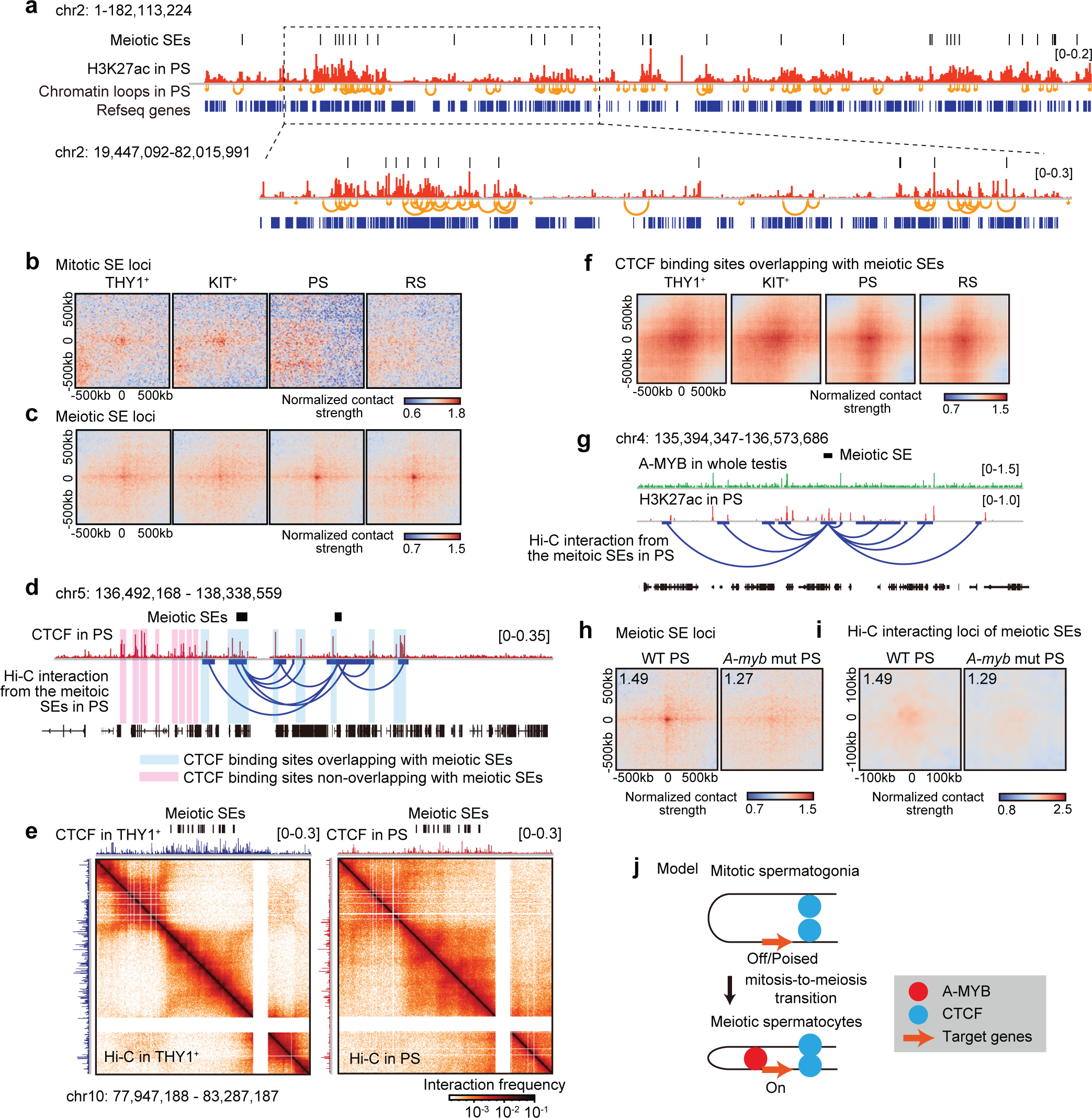
Meiotic super-enhancers are poised with 3D chromatin. **a**, Track view showing meiotic SEs, H3K27ac, and chromatin loops in PS on the entire chromosome 2 (top). Enlargement of the boxed area is shown below. **b**, Pile-up analysis of averaged intersections of mitotic SEs with 500kb paddles. **c**, Pile-up analysis of averaged intersections of meiotic SEs with 500kb paddles. **d**, Track view showing CTCF distribution and Hi-C interactions of the meiotic SEs in PS on a region of chromosome 5. Pink highlights indicate CTCF binding sites that do not overlap with meiotic SEs; blue highlights indicate CTCF binding sites that overlap with meiotic SEs and their loops. **e**, CTCF binding and Hi-C maps of THY1^+^ spermatogonia and PS around meiotic SEs (25kb bins, chr10: 77,947,188-83,287,187). **f**, Pile-up analysis showing average interactions of CTCF binding sites overlapping with meiotic SEs. The pile-up analysis in THY1^+^, KIT^+^, PS, and RS is based on the Hi-C data from each developmental stage and the genomic coordinates of the CTCF binding sites that overlapped with meiotic SEs or their interacting genomic regions. **g**, Track view showing the distributions of A-MYB binding and H3K27ac around meiotic SEs. Hi-C interaction from the meiotic SEs is also shown. **h**, Pile-up analysis showing average interactions of meiotic SEs with 500kb paddles in WT PS and *A-myb* mutant PS. The normalized contact strength in the central pixel is displayed on the top left. **i**, Pile-up analysis showing average interactions of loci that interacted with meiotic SEs based on Hi-C data with 100kb paddles in WT PS and *A-myb* mutant PS. **j**, Model of the predetermination of 3D chromatin at meiotic SE loci via CTCF in mitotic spermatogonia. A-MYB strengthens these 3D contacts in meiotic spermatocytes.

We also defined 107 “mitotic” SE that are specific to THY1^+^ and KIT^+^ spermatogonia and found that mitotic SEs and meiotic SEs exhibited distinct 3D chromatin dynamics during spermatogenesis. At the mitotic SE loci, strong 3D contacts were detected in THY1^+^ and KIT^+^ spermatogonia, which resolved together with the resolution of mitotic SEs in PS and RS (Fig. 5b). On the other hand, at meiotic SE loci, 3D chromatin contacts were detected in THY1^+^ and KIT^+^ spermatogonia prior to the establishment of meiotic SEs and increased upon activation of meiotic SEs in PS (Fig. 5c). Pre-establishment of 3D contacts of meiotic SEs in KIT^+^ was also detected in a representative Hi-C map (Extended Data Fig. 5b). These 3D contacts cover relatively large regions and are distinct from chromatin loops that are detected as local point interactions. To further analyze the 3D chromatin structure around the meiotic SEs, we detected Hi-C interacting loci centered around the meiotic SEs (Extended Data Fig. 5c). A pile-up analysis shows that, consistent with the SE-SE interactions, contact strength of Hi-C interacting loci increase in PS, while modest contacts are already present in mitotic spermatogonia (Extended Data Fig. 5d). These results suggest that 3D contacts at meiotic SEs are preprogrammed in spermatogonia, raising the possibility that meiotic SEs are poised for later activation through 3D chromatin.

Because SEs determine cell type-specific gene expression programs, we next sought to determine how meiotic SEs regulate target genes via 3D chromatin. We detected 611 genes that are overlapping with the genomic region interacting with meiotic SEs. Among these genes, 26 genes are associated with the GO term spermatogenesis, and were largely upregulated in PSs and RSs (Extended Data Fig. 6a and b). In our previous study, we showed that spermatogenesis-related genes adjacent to the meiotic SEs are upregulated during late spermatogenesis^17^. Here, we extend this observation by demonstrating that meiotic SEs also upregulate SE-interacting genes via 3D contacts.

Next, to investigate how meiotic SEs regulate gene expression, we examined the epigenetic states of genes adjacent to meiotic SE and SE-interacting loci. H3K4me2, which is implicated in the poised chromatin state^15^ and associated with poised meiotic SEs^17^, accumulated highly at these genes in KIT^+^ spermatogonia, but decreased in PS upon activation of meiotic SEs (Extended Data Fig. 6c, d). In PS, H3K4me3 and H3K27ac, markers for active promoters and enhancers, are enriched at these loci instead. Together, these results suggest that meiotic SE pre-establish H3K4me2-enriched 3D contacts with target genes in mitotic spermatogonia, and this epigenetic state is reprogrammed to an H3K4me3/H3K27ac-enriched state upon activation of meiotic SEs.

### CTCF predetermines the 3D contacts of the meiotic SEs in spermatogonia

To determine how the 3D contacts of the meiotic SE are predetermined in spermatogonia, we focused on CTCF, which is involved in mediating 3D chromatin contacts via CTCF-CTCF interactions^47, 48^. To examine the relationship between CTCF and meiotic SEs, we extracted 844 CTCF-binding sites that overlapped with meiotic SEs and their interacting genomic regions in PS (Fig. 5d). We also identified 13,690 sites that did not overlapped with those genomic regions (Extended Data Fig. 7a). CTCF enrichment was largely maintained at the 844 CTCF binding sites that overlapped with meiotic SEs from THY1^+^ spermatogonia to PS (Extended Data Fig. 7b), raising the possibility that CTCF-mediated 3D chromatin contacts persist from mitotic spermatogonia to PS. Indeed, a representative Hi-C heatmap shows that 3D chromatin contacts at the meiotic SE loci are pre-established in THY1^+^ spermatogonia and that CTCF is highly enriched at these sites (Fig. 5e). A pile-up analysis confirmed that strong 3D contacts are maintained from THY1^+^ spermatogonia to RS at CTCF binding sites that overlap with meiotic SEs (Fig. 5f). Of note, this feature is specific to SE loci because CTCF-CTCF chromatin loops are largely reprogrammed from THY1^+^ spermatogonia to PS (Fig. 2e). These results demonstrate that CTCF predetermines the 3D contacts of meiotic SEs in spermatogonia, poising them for later activation.

### A-MYB strengthens 3D contacts of meiotic SEs on autosomes

Because A-MYB establishes meiotic SEs^17^, we reasoned that A-MYB strengthens 3D contacts of meiotic SEs in PS. Indeed, a representative track-view shows that A-MYB binds to meiotic SE-interacting loci (Fig. 5g). Specifically, it binds to the promoter regions of 294 of the 611 genes that interact with meiotic SE-interacting loci (48.1%; Extended Data Fig. 7c). To examine the role of A-MYB in the regulation of 3D chromatin at meiotic SEs, we analyzed the Hi-C data of *A-myb* mutant PS. 3D contacts between meiotic SE and the interacting loci were attenuated in the *A-myb* mutant PS compared to wild-type PS, although modest contacts were still observed (Fig. 5h, i). Together with the CTCF analysis, we conclude that there are two regulatory mechanisms for the establishment and maintenance of meiotic SEs: CTCF predetermines the overall 3D contacts of meiotic SEs in mitotic spermatogonia, and A-MYB strengthens these 3D contacts upon activation of meiotic SEs in meiotic spermatocytes (Fig. 5j).

### SCML2 and A-MYB establish the unique 3D chromatin architecture of the meiotic sex chromosomes

During meiosis, sex chromosomes undergo epigenetic programming that is different from autosomes. They are subject to MSCI and form a distinct nuclear compartment called the XY body (also known as the sex body)^11, 18^ (Fig. 1a). After meiosis, the silent XY-chromosomal structure, called postmeiotic sex chromatin (PMSC), persists in haploid spermatids^49^. Previous Hi-C studies demonstrated that meiotic sex chromosomes do not show specific 3D chromatin features^20–24^, supporting the notion that 3D chromatin structures of the sex chromosomes are random throughout a cell population. In our Hi-C data, we confirmed that spermatogonia-type far-cis interactions disappeared from the X chromosome in PS (Fig. 6a). Although there are 26 meiotic SEs on the X chromosome in wild-type PS (Extended Data Fig. 8a), we did not detect loci that interacted with these meiotic SEs (Extended Data Fig. 8b). We did, however, detect an enrichment of short-range interactions (less than 1.5 Mb) on the X chromosome specifically in PS (Extended Data Fig. 8c-e), which might be related to fact that the X chromosome remains unsynapsed during meiotic prophase I.

**Figure 6.**
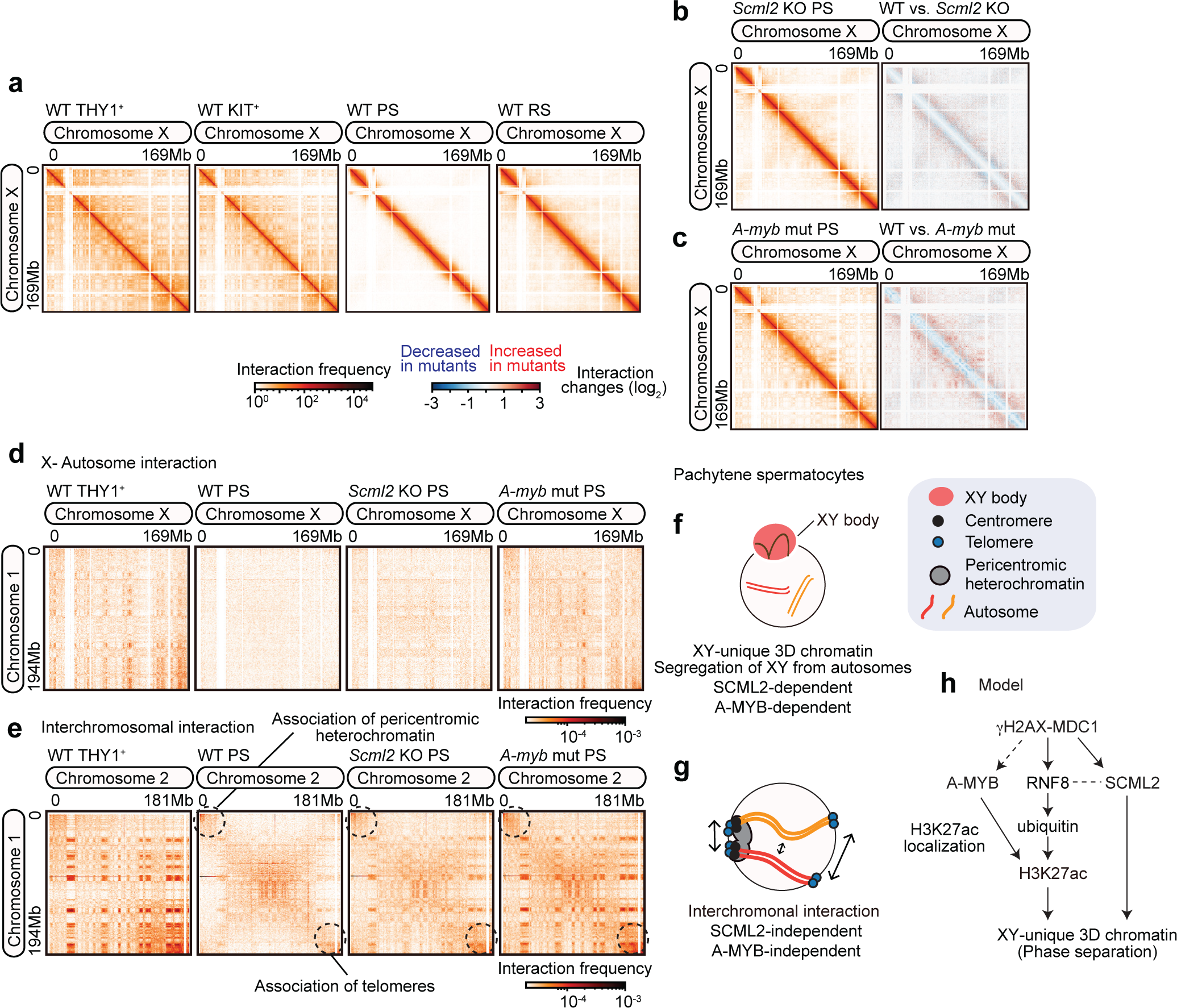
SCML2 and A-MYB establish unique 3D chromatin of the meiotic sex chromosomes. **a**, Hi-C maps of the X chromosome showing normalized Hi-C interaction frequencies (100kb bins) in WT THY1^+^, KIT^+^, PS, and RS. **b**, Heat maps showing normalized Hi-C interaction frequencies (100kb bins, chromosome X) in *Scml2*-KO PS (left). Red and blue Hi-C maps represent a log2 ratio comparison of Hi-C interaction frequencies between wild-type and *Scml2*-KO PS (right). **c**, Heat maps showing normalized Hi-C interaction frequencies (100kb bins, chromosome X) in *A-myb* mutant PS (left). Red and blue Hi-C maps represent a log2 ratio comparison of Hi-C interaction frequencies between wild-type and *A-myb* mutant PS (right). **d**, Heat maps showing normalized Hi-C interchromosomal interactions (250-kb bins, chromosomes 1 and X) for WT THY1*^+^*, WT PS, *Scml2*-KO PS and *A-myb* mutant PS. **e**, Heat maps showing normalized Hi-C interchromosomal interactions (250-kb bins, chromosomes 1 and 2) for WT THY1^+^, WT PS, *Scml2*-KO PS and *A-myb* mutant PS. **f**, Model for the establishment of a unique 3D chromatin in the XY body and segregation of XY from autosomes in PS. **g**, Model of interchromosomal interactions in pachytene spermatocytes. **h**, Schematic of the molecular pathway that establishes a XY-unique 3D chromatin in pachytene spermatocytes.

To determine the mechanisms underlying the unique 3D chromatin organization of the X chromosome, we focused on SCML2 and A-MYB. SCML2 is known to accumulate and function on meiotic sex chromosomes, independently and via a distinct mechanism compared to autosomes^14^. A-MYB is required to establish chromosome-wide accumulation of H3K27ac on the sex chromosome^17^, which facilitates the activation of sex-linked genes in postmeiotic RS ^38^. These functions of SCML2 and A-MYB are regulated downstream of the DNA damage response pathways centered on γH2AX and its binding protein MDC1, which initiate MSCI at the onset of the early pachytene stage^14, 17, 50^. In *Scml2*-KO PS, spermatogonia-type far-cis interactions remain on the X chromosome (Fig. 6b, Extended Data Fig. 8f), and this feature persists in*Scml2*-KO RS (Extended Data Fig. 8h). In *A-myb* mutant PS, spermatogonia-type far-cis interactions also remain on the X chromosome and we observe a plaid pattern of Hi-C signals, which represents the maintenance of spermatogonia-type compartment strengths (Fig. 6c, Extended Data Fig. 8g). These results indicate that both SCML2 and A-MYB are necessary to establish a unique 3D chromatin organization of the X chromosome.

Since meiotic sex chromosome are segregated from autosomes through the formation of the XY body, we examined interchromosomal interactions between the X and autosomes. In wild-type PS, these interchromosomal interactions decreased during spermatogenesis (Fig. 6d, Extended Data Fig. 9). In contrast, in *Scml2*-KO and *A-myb* mutant PS, interchromosomal interactions remained (Fig. 6d, Extended Data Fig. 9), indicating that the segregation of the sex chromosomes form autosomes is dependent on SCML2 and A-MYB (Fig. 6f). Interchromosomal interactions between autosomes remain intact in *Scml2*-KO and *A-myb* mutant PS, including the association of pericentromeric heterochromatin and telomeres as well as the overall association of autosomes (represented by “X” shape signals on Hi-C maps as described previously^51^) (Fig. 6e). Therefore, the role of SCML2 and A-MYB in regulating interchromosomal interactions of meiotic chromosomes is specific to the interaction between the X chromosome and autosomes (Fig. 6g).

In summary, we conclude that SCML2 and A-MYB are required for the establishment of the unique 3D chromatin architecture of the sex chromosomes (Fig. 6h) and the formation of the segregated XY body.

## Discussion

In this study, we determined the high-resolution 3D genome architecture of cell types representative of different stages of spermatogenesis and defined regulatory mechanisms underlying the transition from mitotic spermatogonia to meiotic spermatocytes. We demonstrated that, in spermatogonia, CTCF-mediated 3D contacts at meiotic SE are pre-established. Since meiotic SEs instruct the burst of meiotic gene expression, these poised 3D contacts represent a mechanism to maintain the cellular identity of male germ cells during spermatogenesis. Thus, we show that pre-programming through 3D contacts represents a novel feature of epigenetic priming. Of note, the poised 3D contacts in juvenile spermatogonia reflect the gene expression program of meiotic spermatocytes in adult testis, therefore they are maintained for a long time. Such 3D chromatin-based memories are likely to be prevalent in the germline, as sperm 3D chromatin is preset through histone modifications in late spermatogenesis as well^20^.

Epigenetic priming enables a rapid change in gene expression upon a signaling cue based on a pre-established chromatin state. It has been observed in various biological contexts, including immune cells^52^, neuronal^53^ and cancer development^54^, as well as during spermatogenesis to instruct the gene expression program in late spermatogenesis^15^. Mechanistically, the pre-formation of enhancer-promoter pairs drives transcriptioncal changes upon differentiation in a variety of organisms and cell types^55–57^. Pre-formed enhancer-promoter pairs are associated with paused RNA polymerase^56^ and a recent study showed that meiotic transcription bursts in the male germline are associated with the release of paused RNA polymerase, which is mediated by A-MYB and the testis-specific bromodomain protein BRDT^59^. We propose that at meiotic SEs these mechanisms operate in the context of 3D chromatin. In support of this hypthesis, in somatic cells, SEs are driven by another bromodomain protein, BRD4, as well as Mediator to form liquid-like condensates^60^, thereby providing spatial SE organization. In meiotic spermatocytes, BRDT is expressed in lieu of BRD4^61^. Thus, it is conceivable that preestablished 3D contacts provide venues for A-MYB and BRDT-driven spatial organization of meiotic SEs via phase separation to instruct the burst of meiotic gene expression (Fig. 7a).

**Figure 7.**
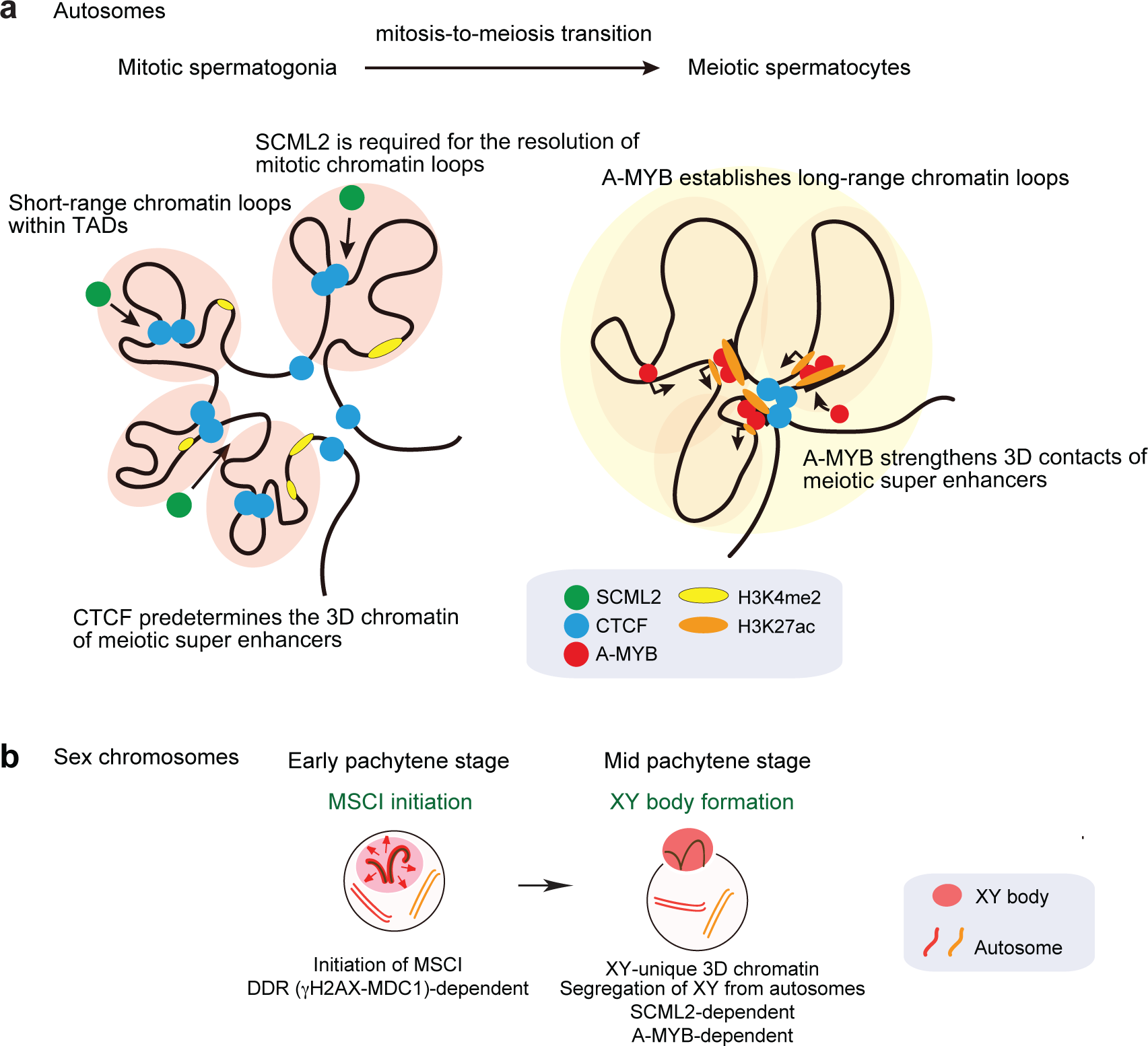
Models of 3D chromatin dynamics and gene regulation on autosomes and sex chromosomes during spermatogenesis. **a**, Model showing the changes in chromosome interactions from mitotic spermatogonia to meiotic spermatocytes on autosomes. **b**, Model of 3D chromatin dynamic on the sex chromosomes. At the onset of MSCI at the early-pachytene stage, DDR initiated MSCI and, subsequently, SCML2 and A-MYB establish unique 3D chromatin of the sex chromosomes and segre-gate the sex chromosomes from autosomes at the mid-pachytene stage.

A key question that remains is the timepoint at which the pre-programmed 3D contacts are established during spermatogenesis. Male germ cells acquire the androgenic epigenome in prospermatogonia (also known as gonocytes) prior to birth^62^. Prospermatogonia are arrested at the G_0_/G_1_ phase of the cell cycle and genome-wide de novo DNA methylation takes place^63, 64^. A recent study has shown that 3D chromatin reprogramming occurs in prospermatogonia^65^. Therefore, it is possible that the 3D contacts necessary for the spermatogenic gene expression program are established in prospermatogonia. Notably, the number of CTCF binding sites is reduced during in vitro differentiation of primordial germ cell-like cells to germline stem cell-like cells^66^, raising the possibility that the androgenic pattern of CTCF binding sites and 3D contacts are reprogrammed in prospermatogonia. CTCFL (also known as BORIS), a paralog of CTCF ^67, 68^, is expressed during spermatogenesis, along with CTCF, and some CTCF-binding sites overlap with CTCFL-binding sites^69^. Hence, CTCFL may be involved in the regulation of CTCF and CTCF-mediated 3D chromatin in spermatogenesis.

When germ cells enter meiosis, cohesin-mediated axial loops are formed along the chromosome axes to promote homolog pairing and recombination. Cytological analyses suggest that an average axial loop length is ∼ hundred(s) kb^46, 70^ in mice, but the average axial loop lengths estimated from contact probability analyses of previous Hi-C studies^24, 51^ is larger (∼ 1 Mb). The average length of stable meiotic chromatin loops detected by our Hi-C analysis is also ∼ 1 Mb (Fig. 1f). Yet, we show that these stable chromatin loops are associated with transcription and are formed around super-enhancers (Fig. 5a), suggesting that they are distinct from axial loops. Further studies using independent approaches are needed to clarify if axial loop structures are distinct from stable chromatin loops.

Finally, our study showed that 3D chromatin organization of the sex chromosomes is regulated by SCML2 and A-MYB. After the initiation of MSCI directed by the DDR pathway at the early pachytene stage, SCML2 localizes on the sex chromosome after the mid-pachytene stage^14^. At that time, A-MYB regulates chromosome-wide spreading of H3K27ac on the sex chromosome^17^, downstream of the DDR factor RNF8^71^. RNF8 interacts with the MSCI initiator, MDCI, which functions as a γH2AX binding protein^50^. Therefore, the DDR pathway coordinates both SCML2 and A-MYB-dependent processes on the sex chromosomes (Fig. 6h). Spermatogonia-type 3D chromatin features are retained in the *Scml2*-KO and *A-myb* mutant PS, suggesting that SCML2 and A-MYB are both required for the establishment of 3D chromatin features of the XY body (Fig. 7b). We suggest that SCML2 and A-MYB may work in concert on the sex chromosomes because SCML2 and RNF8 function together in the regulation of histone ubiquitination on meiotic sex chromosomes^38^. Similar to A-MYB’s function on autosomes, A-MYB could drive a phased separated compartment of the sex chromosomes. Of note, this 3D chromatin feature of the male X chromosome in MSCI is distinct from that of the female inactive X chromosome. The inactivated female X chromosome splits into two mega domains bounded by the *Dxz4* locus and forms long-range loop structures called super loops (>7 Mb)^27^. CTCF binds around the *Dxz4* locus, and this structure is essential for the formation of mega domains and super loops in female cells ^72^. This is quite different from the male X chromosome in meiosis, where short-range interactions are increased, and a domain structure is not clearly visible.

Overall, our results uncover the mechanisms underlying the organization of the meiotic chromatin structure on both autosomes and sex chromosomes and establish that CTCF-mediated pre-programming drives the burst of autosomal gene expression during male meiosis.

## Methods

### Animals and germ cell isolation

Mice were maintained and used according to the guidelines of the Institutional Animal Care and Use Committee (protocol no. IACUC2018-0040) at Cincinnati Children’s Hospital Medical Center. Wild-type C57BL/6J mice, *Scml2*-KO mice^14^ on the C57BL/6J background, and *A-myb*^mut/mut^ (*Mybl1*^repro9^)^25^ on the C57BL/6J background were used for Hi-C analyses. Spermatogonia were isolated from C57BL/6J wild-type aged 6-8 days through magnetic cell-sorting (MACS) as described previously^26^. Pachytene spermatocytes and round spermatids, including *Scml2*-KO PS, Scml2-RS, and *A-myb* mutant PS, were isolated from adult testes through sedimentation velocity at unit gravity as described previously^26, 39, 73^.

### Hi-C library generation and sequencing

To generate and sequence Hi-C libraries, Hi-C was used using the Arima Hi-C kit, according to the manufacturer’s instructions. We used the Arima-Hi-C kit, which enables high-resolution detection of 3D chromatin by using a combination of multiple restriction enzymes. 4×10^5^ to 1×10^6^ cells were used for THY1^+^ and KIT^+^, 3×10^6^ cells were used for WT PS, *Scml2*-KO PS, and *A-myb* mutant PS, and 4×10^6^ cells were used for WT RS and *Scml2*-KO RS for crosslinking. For library preparation, Accel-NGS^®^ 2S Plus DNA Library Kit (Swift Biosciences, Inc. Ann Arbor, MI) was used. All libraries were sequenced on Illumina HiSeq4000 sequencers according to the manufacturer’s instructions.

### Hi-C data mapping

Paired-end .fastq files of Hi-C libraries were aligned and processed using the Juicer package^74^ (version 1.5). In brief, each end of the raw reads was mapped separately to the *Mus musculus* mm10 reference genome, and Hi-C pairs files were created using BWA^75^ (version 0.7.3a). Mapping statistics are summarized in Supplementary Table 1. *.hic* files, a highly compressed binary file was created by Juicer tools pre. Matrix balancing was performed with the cooler software package (version 0.8.11) and visualized using the HiCExplorer^76^ (version 3.6) for use with the application hicPlotMatrix. To generate and visualize interaction frequency heat maps of whole chromosomes, Hi-C matrices at 250-kb resolution were imported to the software package HiCExplorer for use with the application hicPlotMatrix. To aid visual comparisons between the datasets, matrices were natural log transformed. To analyze differential interaction frequencies between samples, the HiCExplorer application hicCompareMatrices was used to generate log2 ratios of interaction frequency matrices between two separate datasets and then visualized by hicPlotMatrix.

### Hi-C: Evaluation of Hi-C biological replicates

Pearson correlation coefficients between Hi-C biological replicates at each stage were obtained by using the hicCorrelate included in HiCExplorer using the cool files binned at 10kb with the parameter ‘--log1p --method pearson’. A range from 10kb to 5Mb was used in the calculations. The reproducibility of the results was confirmed with biological replicates (Extended Data Fig. 1c).

### Hi-C: Contact frequency

Enrichment of Hi-C counts at different genomic ranges/distances to whole chromosomes was calculated for autosomes and X chromosomes respectively using hicPlotDistVsCounts including HiCExplorer with the option ‘--maxdepth 300000000’.

### Hi-C: Identification of chromatin loops

Chromatin loops were called by using the HiCEexplorer for the use with the application hicDetectLoops using cool files binned at 5kb, 10kb and 25kb respectively with the parameter ‘--maxLoopDistance 2000000’. The cool files were generated from hic files created using only reads with a MAPQ score of 30 or higher using the -q 30 option during Juicer tools pre procedure and converted to cool files using the hicConvertFormat included in HiCExplorer and the cooler balance included in cooler. After detecting chromatin loops at each resolution, the loops from each resolution were merged by using the hicMergeLoops included in HiCExplorer with the ‘-r 25000’ option. CTCF, H3K4me3/H3K27ac and H3K27me3-dependent loops were detected using bed files of CTCF (this study), H3K4me3^39^/H3K27ac^17, 38^ or H3K27me3^39^ ChIP-seq datas by the hicValidateLocations with the ‘--method loops --resolution 25000’ option. Juicer tools compare was used to detect common loops between the two types of chromatin loops, with the option ‘-m 25000 0 mm10’. The loop listed as “Common”, “A” or “B” in parent_list of the output data was used as the common loops between the two loops. IGV^77^ (version 2.8.3) was used to visualize chromatin loops in the genomic track view. Genes where the anchor site of the loop overlaps with the TSS region using refTSS_v3.1_mouse_annotation.txt (https://reftss.riken.jp/datafiles/3.1/mouse/gene_annotation/) were identified as a group of genes associated with a specific loop.

### Hi-C: Identification of Topologically Associated Domains (TADs)

TADs were detected by the hicFindTads including HiCExplorer using the cool files binned at 25kb with the parameter ‘--correctForMultipleTesting fdr --minDepth 80000 --maxDepth 800000 --step 40000 minBoundaryDistance 80000 --thresholdComparisons 0.01 --delta 0.01’. TAD boundaries were analyzed by extending 25 kb each upstream and downstream from the TAD boundaries detected by hicFindTads for downstream analysis. CTCF, H3K4me3/H3K27ac and H3K27me3-dependent TAD boundaries were detected using bed files of CTCF, H3K4me3/H3K27ac or H3K27me3 ChIP-seq datas by the hicValidateLocations with the ‘--method tad --resolution 25000’ option. Common TAD boundaries between the two data sets were detected using bedtools intersect, and were assumed to be common if they covered even 1 bp.

### Hi-C: Identification of mitotic and meiotic specific SEs and genomic sites interacting with meiotic SEs

The SE files downloaded from Maezawa et al., 2020^17^ was used with a modified version of the SE file. The SEs detected in THY1^+^ and KIT^+^, excluding those overlapping with the SEs detected in PS and RS, were used as mitotic specific SEs, and conversely, the SEs detected in PS and RS, excluding those detected in THY1^+^ and KIT^+^, were used as meiotic specific SEs in this study. To determine whether PS chromatin loops overlapped with meiotic SE, the anchor stes of PS chromatin loops were added +0.4 Mb upstream and downstream, and it was determined whether this region overlapped with meiotic SE.

The regions interacting with meiotic SEs followed the method described in ‘3.4.6 Identification of super-enhancer-promoter interactions’ by Sakashita et al., 2023^78^. The first step in the analysis is to calculate the quality of each viewpoint (SE locus) using the chicQualityControl program including in HiCExplorer, which considers the sparsity of the Hi-C contact frequency with the ‘--sparsity 0.3’ option. Next, using the bed file containing the filtered viewpoints and the program chicViewpointBackgroundModel, a background model of all given viewpoints is calculated based on the Hi-C contact matrix, with the option to set the range of the background model to 500 kb with the ‘--fixateRange 500000’ option. Using the chicViewpoint program, all interaction points in physical contact with the SE locus are detected based on the background model, ranging up to 500 kb (--range 500000 500000)). Finally, using the chicSignificantInteractions program with the ‘-p 0.05 --range 500000 500000 --loosePValue 0.1’, only significant interaction points (P<0.05 (-p 0.05)) were extracted, which were designated as genomic regions interacting with meiotic specific SEs.

### Hi-C: Pile-up analysis

Pile-up analysis was performed using coolup.py^36^ (version 0.9.5) to visualize the average interaction strength of chromatin loops and TADs. Chromatin loops were analyzed using the cool files binned at 10kb resolution and a bedpe file of the corresponding chromatin loops. TAD domains were analyzed using the cool files binned at 25kb resolution and bed file showing TAD domains with the ‘--rescale --local’ option. TAD boundaries were analyzed using the cool files binned at 25kb resolution and bed file showing TAD boundaries with the ‘--pad 500 --local’ option. For the analysis of interactions between mitotic or meiotic SEs, bed files showing mitotic or meiotic SEs were used. For the pile-up analysis showing the interaction among mitotic or mitosis-specific SEs, each SEs were analyzed using the cool files binned at 10kb resolution and bed file showing TAD boundaries with the ‘--pad 500’ option. For the interaction between CTCFs on meiotic SEs, the overlap regions between meiotic SEs and CTCF binding sites in PS were used for analysis. CTCFs on meiotic SEs were analyzed using the cool files binned at 10kb resolution and bed files showing TAD boundaries with the ‘--pad 500’ option. Piled-up data were visualized by performing plotup.py.

### RNA-seq data analysis

Row RNA-seq reads after trimming by Sickle (https://github.com/najoshi/sickle) (version 1.33) trimmed regions with quality less than 30 and excluded reads that were less than 20 bp. Trimmed sequencing reads were aligned to the *Mus musculus* mm10 reference genome using HISAT2^79^ (version 2.2.1) with default parameters. All unmapped reads and non-uniquely mapped reads were filtered out and then sorted by samtools^80^ (version 1.14) with default parameters. The output bam file was assembled and quantified using StringTie^81^ (version 2.2.1) based on the mouse gene annotation (gencode.vM25.annotation.gtf). Transcripts per million (TPM) value was used for downstream analyses.

Genes associated with *Scml2*-KO PS-specific loops were defined with genes whose TSS regions overlapped with the anchor sites of these loops. Then, only genes with TPM values greater than 1 in any of the cells were extracted and used for analysis. Violinplot was drawn using the R package ggplot2. The log10-transformed values of TPM values+1 were used for statistical analysis and plotting. GO term analysis was performed using the website tool DAVID (https://david.ncifcrf.gov/home.jsp). GO term was visualized by ggplot2 of the R package based on gene number, fold enrichment, and *P* value.

### ChIP-seq data analysis

Cross-linking ChIP-seq was performed for CTCFs of THY1^+^, WT PS, and *Scml2*-KO PS, using the same methods as previously reported^17^. The reproducibility of the results was confirmed with biological replicates (Extended Data Fig. 2a, 3b). Row ChIP-seq reads after trimming by Sickle trimmed regions with quality less than 20 and excluded reads that were less than 20 bp. Trimmed sequencing reads were aligned to the *Mus musculus* mm10 reference genome using Bowtie2^75^ (version 2.4.5) with default parameters. All unmapped reads and non-uniquely mapped reads were filtered out and then sorted by samtools (version 1.14) with default parameters. All unmapped and uniquely mapped reads were filtered out, and sorted by default parameters using samtools, and then ‘MarkDuplicates’ command in Picard tools (version 2.26.9; https://broadinstitute. github.io/picard/) was used to remove PCR duplicates by using the option ‘VALIDATION_STRINGENCY=LENIENT ASSUME_SORTED=true REMOVE_DUPLICATES=true’. After this process, the bam files sorted by samtools again were used for downstream analysis.

To compare biological replicates, Pearson correlation coefficients were calculated and plotted by multiBamSummary bins and plot correlation from deepTools^82^ (version 3.5.1). For visualization of ChIP-seq using IGC, normalized genome coverage tracks based on counts per million mapped reads were generated as bigwig files using bamCoverage function of deepTools with ‘--binSize=5 --normalization CPM’ parameter. Bigwig files were also used for visualization of ChIP-seq data using IGV. Peak calles were identified using MACS2^83^ (version 2.2.7.1). The ngs.plot was used to draw tag density and heat maps for read enrichment within ± 2kb for CTCF and histone modification, ±3kb for A-MYB analysis at meiotic SEs interacting sites, and ± 5kb for A-MYB analysis at anchor sites of PS chromatin loops^84^. A-MYB binding genes were extracted using the online website GREAT (version 4.0.4; http://great.stanford.edu/public/html/) for genes with TSS in the peak +-2 kb region of A-MYB ChIP-seq.

### Statistics

Statistical methods and *P* values for each plot are listed in the figure legends and/or in the Methods. For all experiments, no statistical methods were used to predetermine sample size. Experiments were not randomized, and investigators were not blinded to allocation during experiments and outcome assessments.

### Data availability

Hi-C and CTCF ChIP-seq datasets were deposited in the Gene Expression Omnibus under accession no. GSE244681. All other next-generation sequencing datasets used in this study are publicly available. RNA-seq data from THY1^+^ spermatogonia, PS and RS were downloaded from the GEO (accession no. GSE55060). ChIP-seq data for H3K4me2, H3K4me2 and H3K27me3 and RNA-seq data from KIT^+^ spermatogonia were downloaded from the GEO (GSE89502). ChIP-seq data for H3K27ac in WT PS were downloaded from the GEO (GSE107398). H3K27ac in THY1^+^ and KIT^+^ spermatogonia and input for CTCF ChIP-seq were downloaded from the GEO (GSE130652). A-MYB ChIP seq in whole testis was downloaded from GEO (GSE44588). Source data are provided in this paper.

### Code availability

Source code for all software and tools used in this study, with documentation, examples, and additional information, is available at the URLs listed above.

## Acknowledgments

We thank members of the Namekawa lab, Kris Alavattam, Brad Cairns, and Chongil Yi for the discussion, Xin Li for sharing A-myb mutant mice, and Artem Barski for sharing the reagents.

## Author contributions

Y.K. and S.H.N. designed the study. K.T. and S.M performed experiments. Y.K. performed the computational analyses. Y.M. contributed to the computational analyses. A.S. contributed to developing a computational tool. Y.K., N.K, and S.H.N interpreted the computational analyses. Y.K. and S.H.N. wrote the manuscript with critical feedback from all other authors. S.H.N. supervised the project.

