## Extended Figs for "CTCF-mediated 3D chromatin predetermines the gene expression program in the male germline"

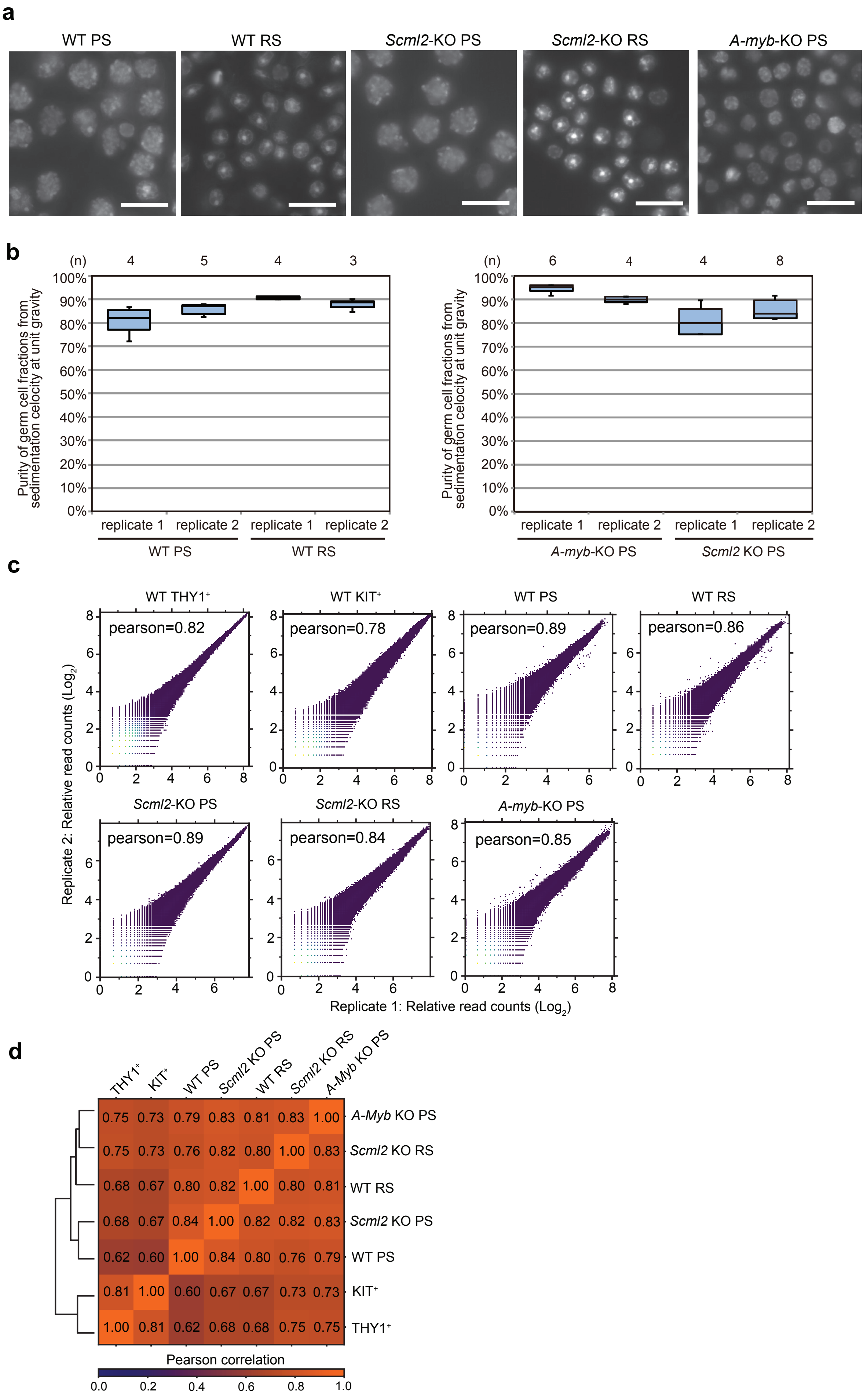
a
WT PS
WT RS
Scml2-KO PS
Scml2-KO RS
A-myb-KO PS
b
(n) 4 5 4 3
Purity of germ cell fractions from sedimentation celocity at unit gravity
100% 90% 80% 70% 60% 50% 40% 30% 20% 10% 0%
replicate 1 replicate 2 replicate 1 replicate 2
WT PS
WT RS
(n) 6 4 4 8
A-myb-KO PS
Scml2 KO PS
c
WT THY1+
pearson=0.82
WT KIT+
pearson=0.78
WT PS
pearson=0.89
WT RS
pearson=0.86
Scml2-KO PS
pearson=0.89
Scml2-KO RS
pearson=0.84
A-myb-KO PS
pearson=0.85
Replicate 2: Relative read counts (Log2)
Replicate 1: Relative read counts (Log2)
d
THY1+ KIT+ WT PS Scml2 KO PS WT RS Scml2 KO RS A-Myb KO PS
0.75 0.73 0.79 0.83 0.81 0.83 1.00 A-Myb KO PS
0.75 0.73 0.76 0.82 0.80 1.00 0.83 Scml2 KO RS
0.68 0.67 0.80 0.82 1.00 0.80 0.81 WT RS
0.68 0.67 0.84 1.00 0.82 0.82 0.83 Scml2 KO PS
0.62 0.60 1.00 0.84 0.80 0.76 0.79 WT PS
0.81 1.00 0.60 0.67 0.67 0.73 0.73 KIT+
1.00 0.81 0.62 0.68 0.68 0.75 0.75 THY1+
Pearson correlation
0.0 0.2 0.4 0.6 0.8 1.0

**Extended Data Figure 1. Germ cell isolation for Hi-C libraries and evaluation of biological replicates.**

**a**, Representative DAPI-stained images of isolated cell fractions for wild-type (WT) pachytene spermatocytes (PS), round spermatids (RS), *Scml2*-KO PS, *Scml2*-KO RS, and *A-Myb* mutant PSs. Scale bars: 20µm.

**b**, Box plots showing the percent purity of cell fractions obtained from sedimentation velocity at unit gravity for each biological replicate (replicates 1 and 2) for the WT PSs, WT RSs, *A-Myb* mutant PSs, and *Scml2*-KO PSs Hi-C library. Numbers (n) along the top indicate the numbers of fractions used to prepare the corresponding library replicates below. The box indicates the 25th, median and 75th percentiles, whiskers the 10th to 90th percentiles.

**c**, Scatter plots showing the Pearson correlation between the two biological replicates for the Hi-C dataset. Pearson correlation values are computed by HiCExplorer using the two-sided Pearson method.

**d**, Heatmap showing the pairwise correlation coefficients among each cell type. Two biological replicates were merged for this analysis.

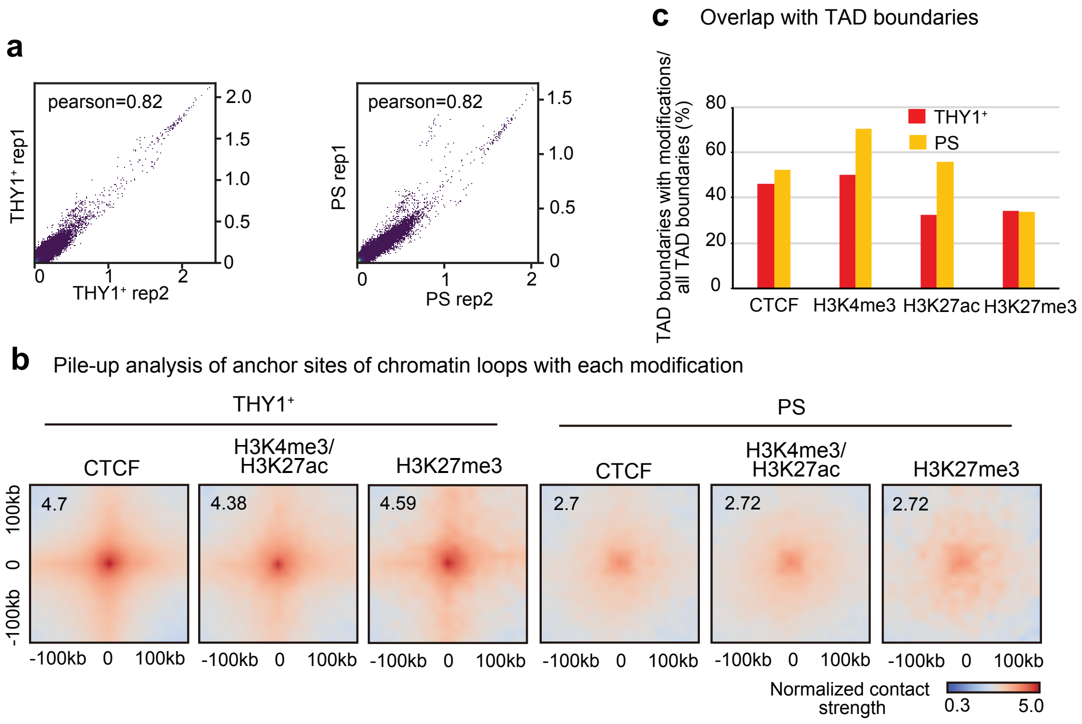


**Extended Data Figure 2. CTCF distribution and chromatin loops at THY1$^+$ and PS.**

**a**, Scatter plots showing the Pearson correlation between 2 biological replicates for CTCF ChIP-seq data in THY1$^+$ and PS. Pearson correlation values are computed by deepTools using the two-sided Pearson method.

**b**, Pile up analysis of CTCF-specific, H3K4me3/H3K27ac-specific, H3K27me3-specific chromatin loops in THY1$^+$ and PS with 100kb padding. The normalized contact strength in the central pixel is displayed on the top left.

**c**, Ratio of accumulation of CTCF, H3K4me3/H3K27ac, or H3K27me3 at TAD boundaries in THY1$^+$ and PS.

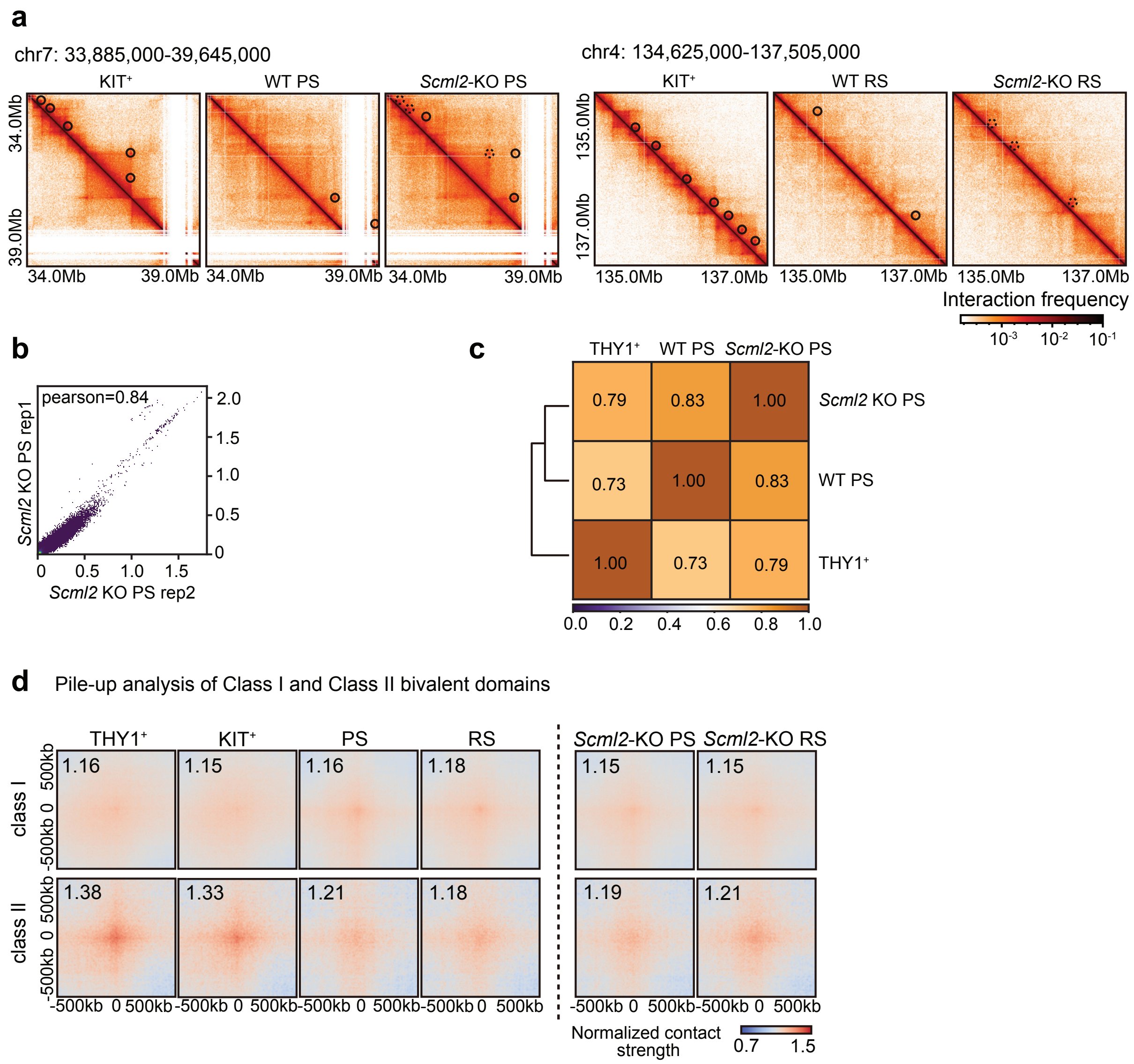


**Extended Data Figure 3. Hi-C analysis of *Scml2*-KO PS and RS.**

**a**, Hi-C maps showing chromatin loops (25kb bins, chr7: 33,885,000-39,645,000 and chr4: 134,625,000-137,505,000) in KIT+, wildtype (WT) PS, and *Scml2*-KO PS. Chromatin loops are indicated by black circles. The dotted circles in the Hi-C map of *Scml2*-KO cells indicate the loops also detected in KIT+.

**b**, Scatter plots showing the Pearson correlation between 2 biological replicates of CTCF ChIP-seq data in *Scml2*-KO PS.

**c**, Heatmap showing the pairwise correlation coefficients for CTCF ChIP-seq in WT THY1+, WT PS, and *Scml2*-KO PS.

**d**, Pile-up analysis showing averaged intersections of class I or class II bivalent domains with 500kb paddles. The genomic coordinates of Class I or Class II bivalent domains are downloaded from [39]. The normalized contact strength in the central pixel is displayed on the top left.

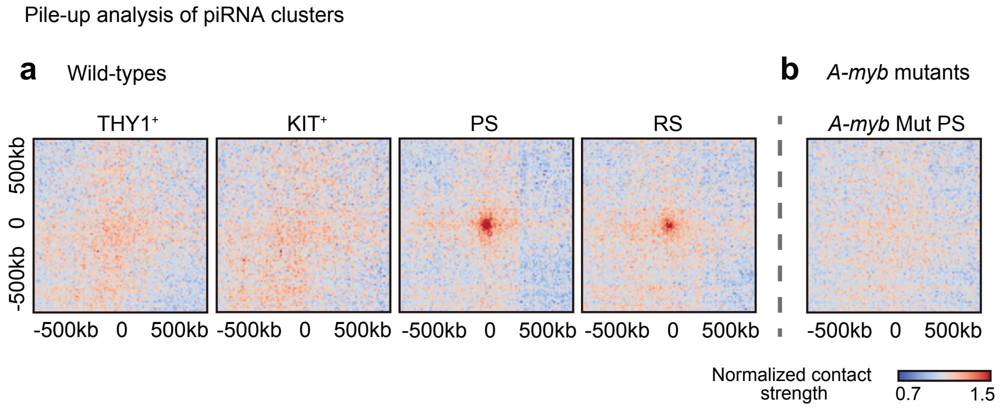


**Extended Data Figure 4. Hi-C analysis of pachytene piRNA loci.**

**a**, **b**, Pile-up analysis showing averaged intersections of pachytene piRNAs with 500kb paddles in wild-type cells (**a**) and *A-myb* mutant PS (**b**). The genomic coordinates of pachytene piRNA loci are downloaded from [40].

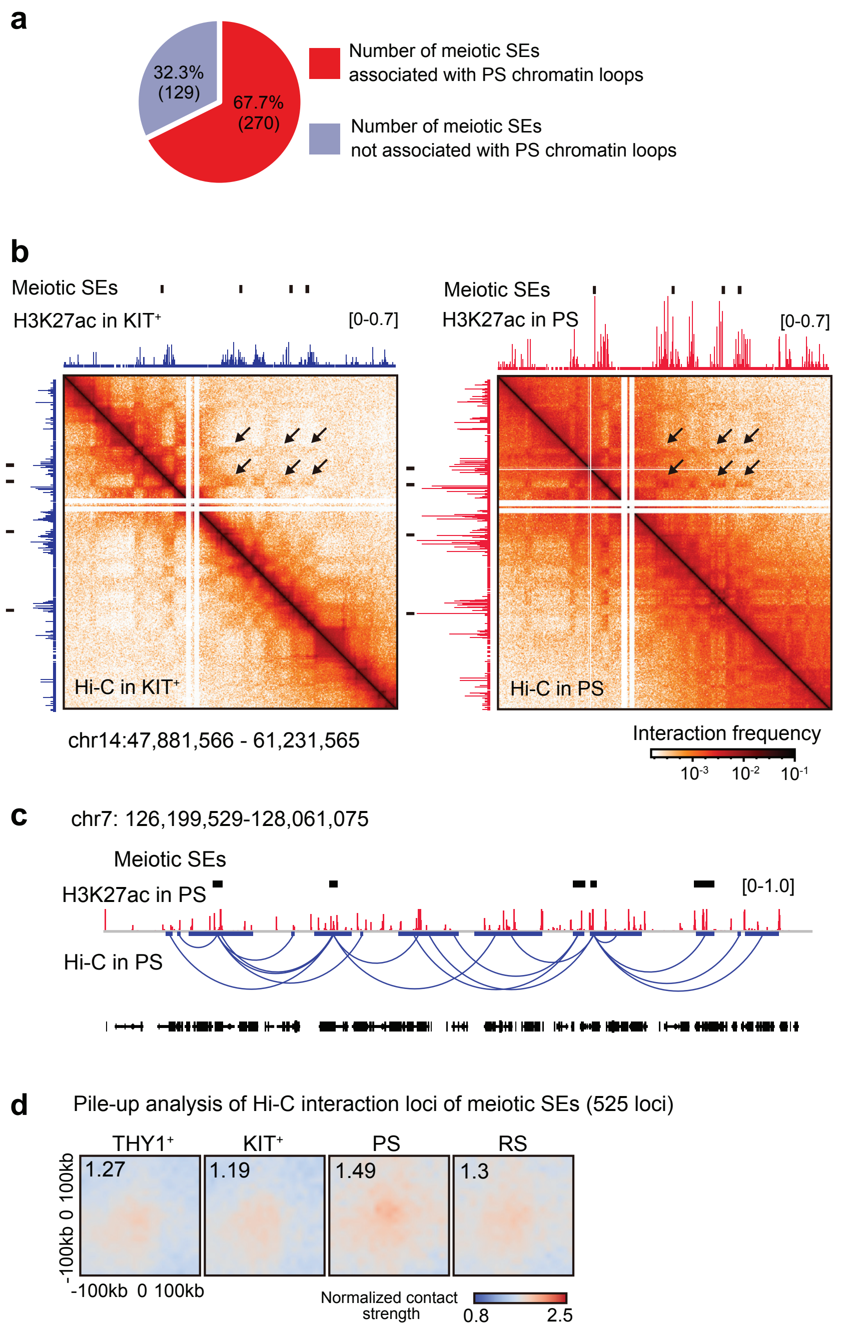


**Extended Data Figure 5. Hi-C analysis of meiotic SEs.**

**a**, Pie chart showing the percentage of meiotic SEs located around chromatin loops. The anchor site and 0.4 Mb upstream and downstream of the anchor site were defined as the genomic region associated with the PS chromatin loop. X and Y chromosomes were excluded from this analysis.

**b**, Heat maps showing normalized Hi-C interaction frequencies (10kb bins, chromosome 14,47,881,566-61,231,565bp) in KIT$^{+}$ and PS. Black arrows highlight meiotic SE-to-SE interactions. ChIP-seq data of H3K27ac in KIT+ and PS are displayed blue and red, respectively.

**c**, Track view showing meiotic SEs (black bars, top), H3K27ac in PS (red), and Hi-C interaction from the meiotic SEs (blue curves).

**d**, Pile-up analysis of Hi-C interaction from the meiotic SEs in each cell type with 100kb padding. The normalized contact strength in the central pixel is displayed on the top left.

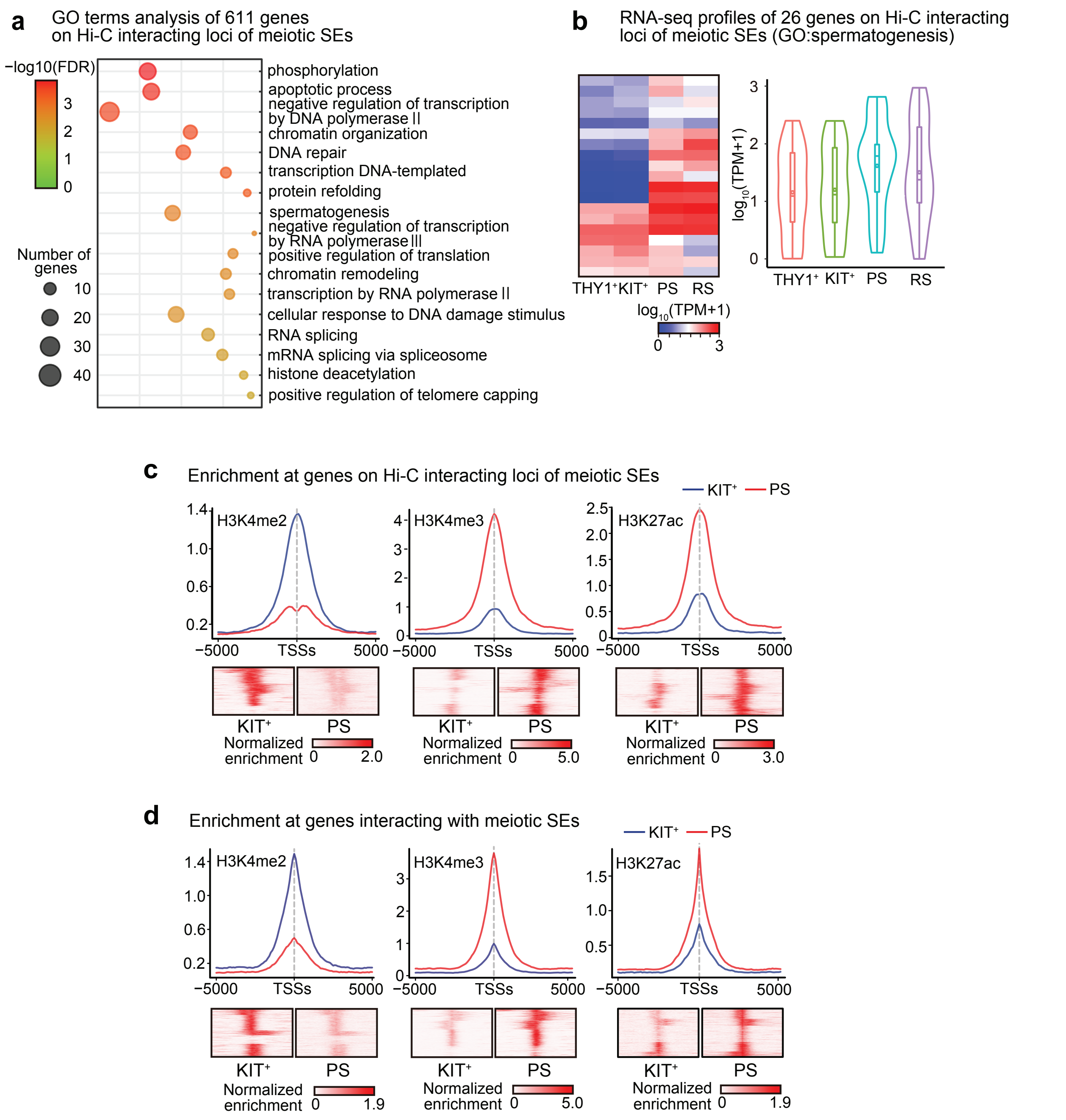


**Extended Data Figure 6. Meiotic SE interacting genes.**

**a**, Gene ontology (GO) analysis of genes interacting with meiotic SEs.

**b**, A heatmap and violin plots showing gene expression of 26 'spermatogenesis' genes that showed Hi-C interaction with meiotic SEs during spermatogenesis. The box in the violin plots indicates the 25th, median, and 75th percentiles, and the dot in the box indicates the mean. All genes that show at least Transcripts per million (TPM 4) at one developmental stage are shown.

**c**, **d**, Average tag densities and heatmaps for H3K4me2, H3K4me3, and H3K27ac ChIP-seq enrichment 5,000bp upstream and downstream of TSSs. Genes adjacent to meiotic SEs (**c**) and genes interacting with meiotic SEs (**d**) are analyzed.

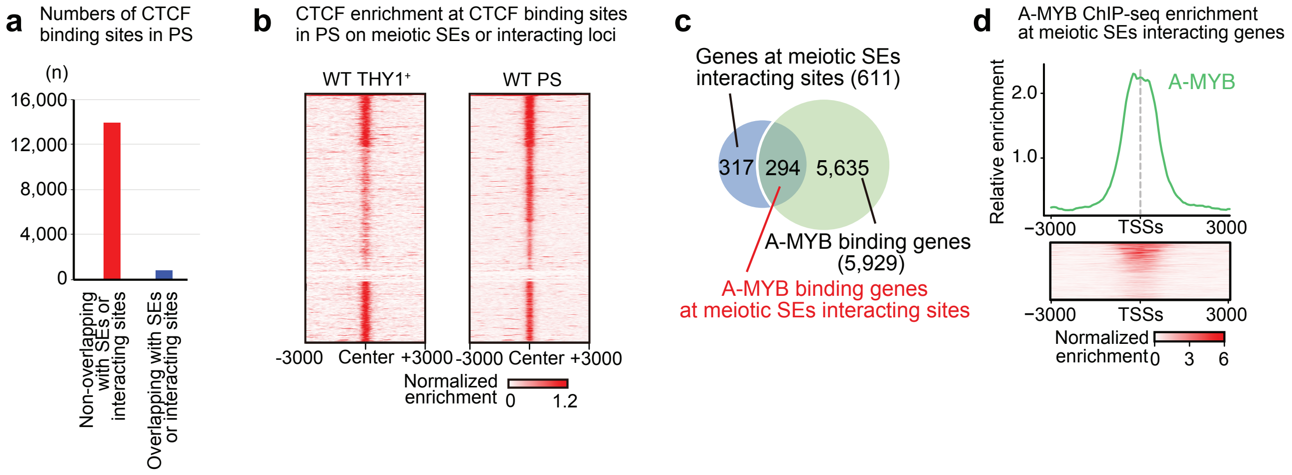


**Extended Data Figure 7. Meiotic SEs poising with 3D chromatin**

**a**, Number of CTCF binding sites in PS that overlap or do not overlap with meiotic SE and meiotic SE interacting loci.

**b**, Heatmap showing CTCF enrichment in $THY1^{+}$ and PS at CTCF binding sites on meiotic SEs in PS.

**c**, Venn diagram showing the intersection between all the genes at meiotic SE interacting genomic regions (blue) and A-MYB binding genes (green).

**d**, Average tag density and heatmap for A-MYB ChIP-seq enrichment 3,000bp upstream and downstream of TSSs of meiotic SE interacting genes.

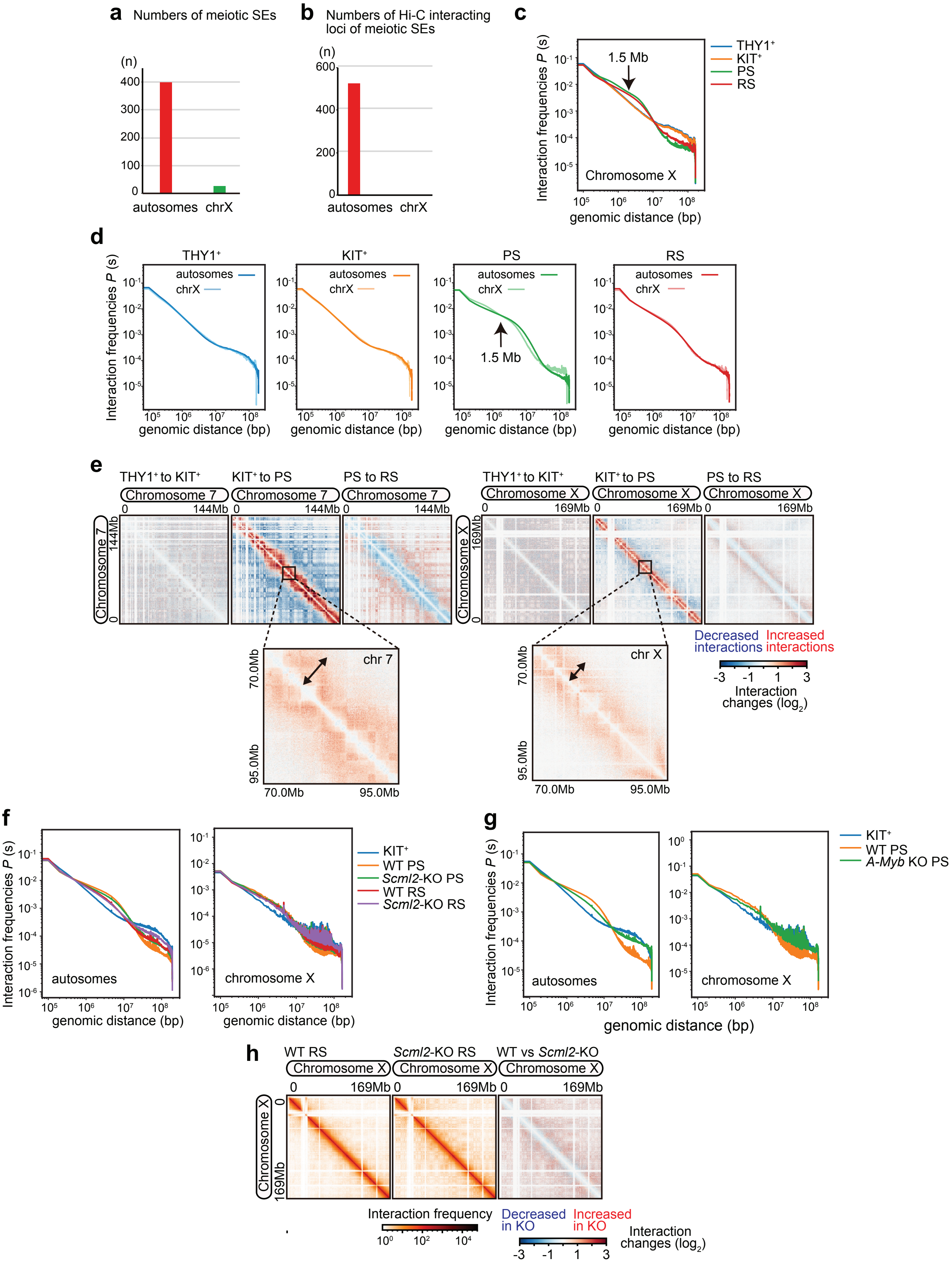

a Numbers of meiotic SEs
(n)
400
300
200
100
0
autosomes chrX
b Numbers of Hi-C interacting loci of meiotic SEs
(n)
600
400
200
0
autosomes chrX
c
Interaction frequencies P (s)
1.5 Mb
Chromosome X
genomic distance (bp)
THY1+
KIT+
PS
RS
d
THY1+
KIT+
PS
RS
autosomes
chrX
1.5 Mb
Interaction frequencies P (s)
genomic distance (bp)
e
THY1+ to KIT+
KIT+ to PS
PS to RS
Chromosome 7
144Mb
Chromosome X
169Mb
chr 7
chr X
70.0Mb
95.0Mb
Decreased interactions
Increased interactions
-3 -1 1 3
Interaction changes (log2)
f
KIT+
WT PS
Scml2-KO PS
WT RS
Scml2-KO RS
autosomes
chromosome X
Interaction frequencies P (s)
genomic distance (bp)
g
KIT+
WT PS
A-Myb KO PS
autosomes
chromosome X
Interaction frequencies P (s)
genomic distance (bp)
h
WT RS
Scml2-KO RS
WT vs Scml2-KO
Chromosome X
0
169Mb
Interaction frequency
Decreased in KO
Increased in KO
Interaction changes (log2)
-3 -1 1 3

**Extended Data Figure 8. Characteristics of chromosome interactions in the X Chromosome.**
**a**, Numbers of meiotic SEs on autosomes and the X chromosome.
**b**, Numbers of Hi-C interacting loci of meiotic SEs on autosomes and on the X chromosome.
**c**, Hi-C interaction frequency probabilities P of the X chromosome stratified by genomic distance s for each cell type shown (100kb bins).
**d**, Comparison of Hi-C interaction frequency probabilities P between autosomes and the X chromosome for each cell type (100kb bins).
**e**, Log2 ratio comparisons of the Hi-C interaction frequencies (100kb bins, chromosome 7 and X) for successive cell types. Arrows indicate the distance of chromosome interactions. 10kb bins normalized Hi-C matrices were used for the zoom-in.
**f**, Hi-C interaction frequency probabilities P stratified by genomic distance s for $KIT^+$, WT PS, *Scml2*-KO PS, WT RS, and *Scml2*-KO RS shown (100kb bins). Autosomes and the X chromosome are shown separately.
**g**, Hi-C interaction frequency probabilities P stratified by genomic distance s for $KIT^+$, WT PS, and *A-myb* mutant PS (100kb bins). Autosomes and the X chromosome are shown separately.
**h**, Heat maps showing normalized Hi-C interaction frequencies (100kb bins, chromosome X) in WT RS, *Scml2*-KO RS. Red and blue Hi-C maps represent a log2 ratio comparison of Hi-C interaction frequencies between wild-type and *Scml2*-KO RS.

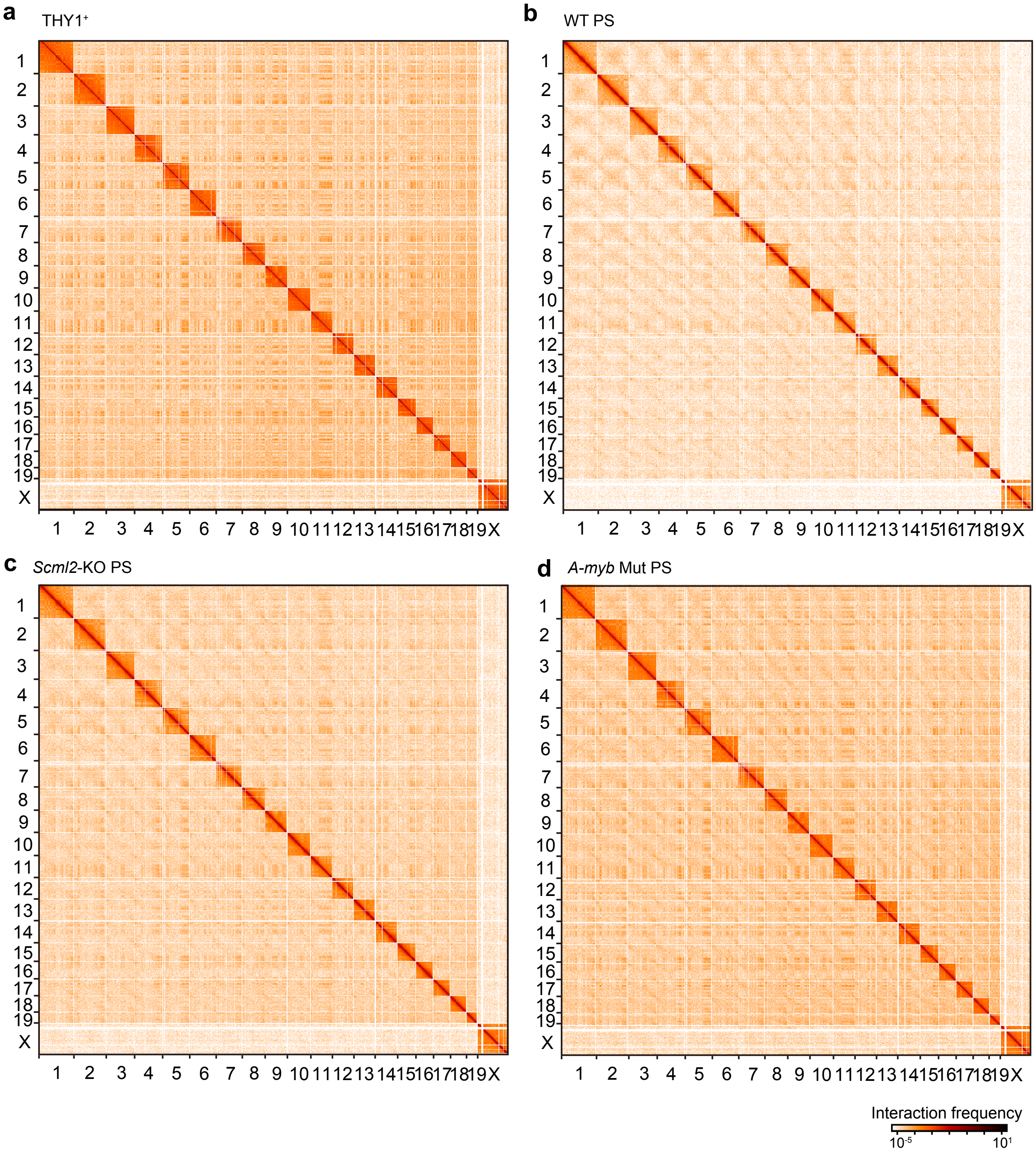


**Extended Data Figure 9. Interchromosomal interactions.**

**a-d**, Heat map showing genome-wide normalized Hi-C interaction frequencies (250-kb bins) in THY1+ (**a**), WT PS (**b**), *Scml2*-KO PS (**c**) and *A-Myb* mutant PS (**d**).
